# Proteome-scale induced proximity screens reveal highly potent protein degraders and stabilizers

**DOI:** 10.1101/2022.08.15.503206

**Authors:** Juline Poirson, Akashdeep Dhillon, Hanna Cho, Mandy Hiu Yi Lam, Nader Alerasool, Jessica Lacoste, Lamisa Mizan, Mikko Taipale

## Abstract

Targeted protein degradation and stabilization are promising therapeutic modalities due to their potency and versatility. However, only few E3 ligases and deubiquitinases have been harnessed for this purpose. Moreover, there may be other protein classes that could be exploited for protein stabilization or degradation. Here, we used a proteome-scale platform to identify hundreds of human proteins that can promote the degradation or stabilization of a target protein in a proximity-dependent manner. This allowed us to comprehensively compare the activities of human E3s and deubiquitinases, characterize non-canonical protein degraders and stabilizers, and establish that effectors have vastly different activities against diverse targets. Notably, the top degraders were more potent against multiple therapeutically relevant targets than the currently used E3s CBRN and VHL. Our study provides a functional catalogue of effectors for targeted protein degradation and stabilization and highlights the potential of induced proximity screens for discovery of novel proximity-dependent protein modulators.

## INTRODUCTION

Modulation of target protein stability with small molecules has emerged as one of the most promising innovations in drug discovery (Békés et al., 2022; Deshaies, 2020; Gerry and Schreiber, 2020). In targeted protein degradation (TPD), an interaction between a target protein and an E3 ubiquitin ligase is induced with a small molecule, leading to target ubiquitination and proteasome-dependent degradation. This can be brought about by either heterobifunctional molecules known as Proteolysis-Targeting Chimeras (PROTACs) or molecular glues. PROTACs consist of two covalently linked protein-binding moieties, one of which binds the target while the other recruits the E3 ligase. Molecular glue degraders, such as thalidomide and its analogs, directly contact both the E3 ligase and the target protein, inducing a non-native or enhancing a weak protein-protein interaction that results in degradation of the neo-substrate. Analogous to TPD, targeted protein stabilization (TPS) with Deubiquitinase-Targeting Chimeras (DUBTACs) was recently developed (Henning et al., 2022a). In this approach, a deubiquitinase (DUB) is recruited to the target with a heterobifunctional molecule, inducing deubiquitination and subsequent stabilization of the target.

Modulating protein levels via TPD or TPS has several advantages over traditional small molecule inhibitors or activators. First, TPS and TPD molecules can function sub-stoichiometrically, since one molecule can induce the degradation or stabilization of multiple target protein molecules, respectively. Moreover, they expand the druggable proteome, as they can target any druggable pocket of a protein rather than just functionally relevant sites (Schneider et al., 2021). This is further potentiated by the ability of molecular glues to target surfaces that have been traditionally considered undruggable (Bussiere et al., 2020; Fischer et al., 2014). Finally, experiments with promiscuous inhibitors made into PROTACs have revealed that PROTACs provide an additional layer of target specificity, which is particularly relevant for conserved protein families (such as kinases) with high likelihood of off-target toxicity (Békés et al., 2022; Bondeson et al., 2018; Donovan et al., 2020)

Despite the high therapeutic potential of TPD and TPS, a major barrier in their development is that only few E3 ligases – and a single DUB – have been successfully harnessed for the approach. For example, the human genome encodes over 600 E3 ligases, but the majority of current degraders in clinical development use only two E3s: the thalidomide target Cereblon (CRBN) and the tumor suppressor VHL (Békés et al., 2022). But compounds exploiting these E3s do not work with every target or targeting moiety, significantly hindering the development of novel degraders. Moreover, cellular resistance to degraders can arise via mutations in the hijacked E3 ligase or other components of the ubiquitin-proteasome system (UPS), which would lead to cross-resistance to any molecules harnessing the same E3 complexes (Hanzl et al., 2022; Lu et al., 2018; Mayor-Ruiz et al., 2019; Ottis et al., 2019; Shirasaki et al., 2021; Sievers et al., 2018; Zhang et al., 2019a). Finally, E3 ligases and DUBs with restricted expression profiles could enable tissue-specific modulation of target protein abundance and alleviate potential compound toxicity. Thus, identifying novel effectors for TPD and TPS is one of the key challenges in the development of next generation degraders and stabilizers (Békés et al., 2022).

Recent work has uncovered a handful of additional proteins amenable to TPD or TPS either by chemical proof-of-concept molecules or as protein fusions (Bery et al., 2019; Bussiere et al., 2020; Henning et al., 2022b; Kanner et al., 2020; Lim et al., 2020; Spradlin et al., 2019; Ward et al., 2019; Wei et al., 2021; Zhang et al., 2019b), but most human E3s and DUBs have not been functionally assessed for this purpose. Moreover, the human genome likely encodes other effectors beyond the usual suspects. For example, annotation of canonical E3s and DUBs is based on their characteristic protein domains rather than their molecular function, although other protein classes can also act as substrate adaptors for protein degradation or stabilization (Yoshida et al., 2013; Zeqiraj et al., 2015; Zha et al., 2015). In addition, the development of autophagy-targeting chimeras (AUTACs) and lysosome-targeting chimeras (LYTACs) has demonstrated that other degradative pathways could also be exploited for TPD (Banik et al., 2020; Takahashi et al., 2019), but which proteins in the human proteome could be hijacked for these alternative pathways remains unknown.

Here, we report a functional survey of human proteins that can degrade or stabilize a non-physiological substrate protein in a proximity-dependent manner, generating a comprehensive catalogue of effectors that could be potentially harnessed for targeted protein degradation or stabilization.

## RESULTS

### A platform for proximity-dependent regulation of protein stability

We employed two approaches to discover proximity-dependent effectors of protein stability at proteome scale. The methods rely on co-expression of a target (sensor) and an effector protein in cells with tags that allow for induction of their interaction. As the target, we employed an EGFP-ABI1 fusion protein stably expressed in 293T cells from a bicistronic vector that also expressed TagBFP translated from an internal ribosome entry site (IRES)(**Figure 1A**). In this cell line, we expressed effectors fused to either the abscisic acid (ABA) receptor PYL1, a domain that binds ABI1 in the presence of ABA (Alerasool et al., 2022; Liang et al., 2011), or to vhhGFP, a nanobody that binds to EGFP with high affinity (Caussinus et al., 2012; Saerens et al., 2005). With this design, effector proteins can be brought to the proximity of EGFP-ABI1 either constitutively (vhhGFP) or by chemical dimerization (PYL1)(**Figure 1A**). The levels of EGFP-ABI1 can then be followed by flow cytometry, whereas TagBFP (which does not bind vhhGFP) is used for normalization.

**Figure 1:**
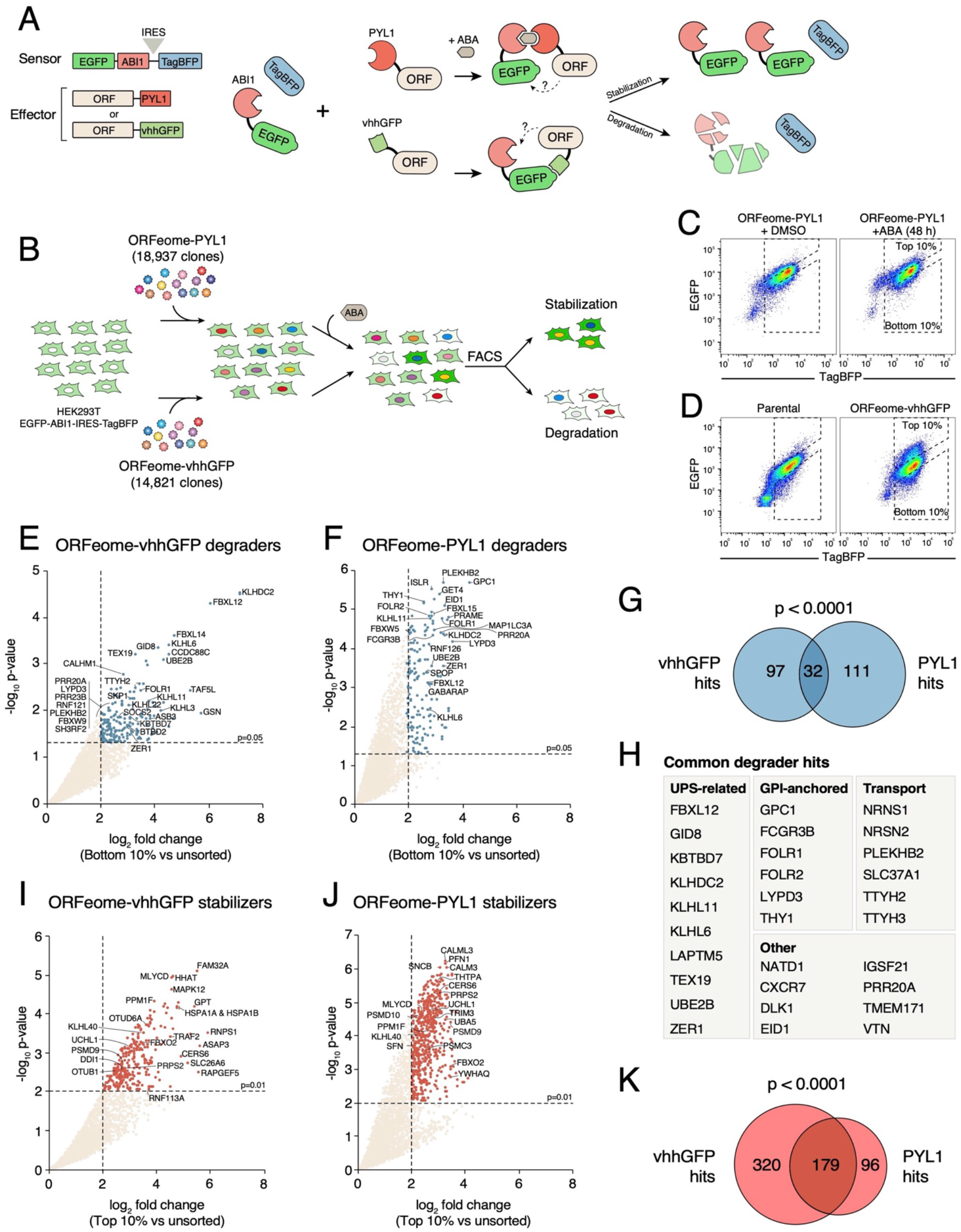
Pooled ORFeome screens for protein stability regulators. (**A**) Schematic of the two approaches used to discover proximity-dependent effectors of protein stability. The method relies on co-expression of an EGFP-ABI1-IRES-TagBFP reporter with effectors fused to tags that bring two proteins together either in a constitutive (vhhGFP/EGFP) or chemically inducible (ABI1/PYL1 + ABA) manner. The levels of EGFP-ABI1 can be followed by flow cytometry and compared to the TagBFP control. (**B**) Outline of the pooled ORFeome screens for protein stability regulators. (**C**) Top 10% and bottom 10% of reporter cells infected with the ORFeome-PYL1 library were sorted in the presence of abscisic acid. (**D**) Similar FACS gates were used for the sorting of the ORFeome-vhhGFP infected cells. (**E**) and (**F**) Enrichment of ORFs in the low EGFP pool infected with the ORFeome-vhhGFP (E) or ORFeome-PYL1 (F) compared to unsorted cells. Significantly enriched ORFs are shown in blue. (**G**) Overlap of genes significantly enriched in the ORFeome-vhhGFP and ORFeome-PYL1 degradation screens. (**H**) List of ORFs identified in both degradation screens. (**I**) and (**J**) Enrichment of ORFs in the high EGFP pool infected with the ORFeome-vhhGFP (I) or ORFeome-PYL1 (J) compared to unsorted cells. Significantly enriched ORFs are shown in red. (**K**) Overlap of genes significantly enriched in the ORFeome-vhhGFP and ORFeome-PYL1 degradation screens.

As a proof of principle, we assessed the effect of two E3 ligases, CRBN and SPOP, on EGFP stability. CRBN is the primary target of thalidomide and commonly used as the effector E3 for molecular glues and PROTACs (Ito et al., 2010; Winter et al., 2015), whereas SPOP has been shown to potently degrade target proteins when fused to vhhGFP (Shin et al., 2015). Both CRBN-PYL1 and SPOP-PYL1 degraded the reporter in an ABA-dependent manner, although SPOP was more potent, in particular when the substrate-binding MATH domain was deleted (**Figure S1A**). Degradation occurred in a time- and dose-dependent manner (**Figure S1B**). The results were similar with vhhGFP fusions (**Figure S1C**). In contrast, we did not observe any target degradation with Nanoluc-PYL1, whereas Nanoluc-vhhGFP expression led to slight stabilization of the reporter (**Figures S1A** and **S1C**).

### ORFeome-wide induced proximity screen for protein stability effectors

To scale up from individual clones, the EGFP-ABI1 reporter cell line was then transduced at low multiplicity of infection with a pooled human ORFeome library tagged with either vhhGFP or PYL1 (Alerasool et al., 2022)(**Figure 1B**). The reporter cells infected with the vhhGFP library or the PYL1 library (treated with ABA) were sorted by flow cytometry based on EGFP/TagBFP intensity ratio in triplicate and duplicate, respectively. The ORFs present in the top and bottom 10% EGFP/TagBFP ratio and the unsorted bulk population were sequenced to assess the relative enrichment of ORFs in the sorted populations (**Figures 1C** and **1D**).

Using a 5% false discovery rate cut-off and at least 4-fold change in normalized read counts between bottom 10% sorted population and unsorted cells, we identified 162 and 179 putative degradation effectors in the vhhGFP and PYL1 screens, respectively (**Figures 1E** and **1F** and **Table S1**). There was a highly significant overlap between the hits identified in the two screens (**Figure 1G**), establishing that many effectors are capable to induce target degradation when brought in proximity either constitutively or by chemical dimerization. Furthermore, confirming the biological relevance of our approach, analysis of the top degradation effectors revealed significant enrichment in regulators of ubiquitination and in several domains characteristic of E3 ligases (**Figures S2A-S2C**). The E3 ligases we identified in the screens represented multiple families, including individual RING E3 ligases (e.g., RNF121) and components of multiprotein E3 complexes such as the CTLH complex (GID8)(Lampert et al., 2018), the BAG6 membrane protein quality control complex (GET4 and RNF126)(Mock et al., 2017), CUL1 Cullin-RING ligases (CRLs; FBXL12 and FBXL15), CUL2 CRLs (ASB3 and PRAME), and CUL3 CRLs (KLHDC2 and KLHL6)(**Figures 1E** and **1F** and **Table S1**). Of note, VHL was not recovered in either screen, suggesting that it is not a particularly robust effector in this context. (CRBN was not part of the ORFeome library.) Degradation inducers also included proteins involved in autophagy, such as GABARAP, MAP1LC3A, and ATG14. Furthermore, several other hits, although not E3 ligases, have been implicated in protein degradation, such as LAPTM5, NDFIP2, and TEX19 (Kawai et al., 2014; MacLennan et al., 2017; Mund and Pelham, 2010). Interestingly, however, many top hits were not related to the ubiquitin-proteasome system, suggesting that the proteome has a previously uncharacterized cache of proximity-dependent degraders (**Figure 1H**). In particular, we identified many glycosylphosphatidylinositol (GPI) anchored proteins and other plasma membrane proteins as hits. Characterization of some of these proteins is described below.

On the stabilization effector side, using a 1% FDR cut-off and at least 4-fold change in normalized read counts, we identified 544 and 297 putative effectors with the vhhGFP and PYL1 screens, respectively (**Figures 1I** and **1J** and **Table S1**). Again, there was a highly significant overlap between the two screens (**Figure 1K** and **Table S1**). Among the overlapping hits were two deubiquitinating enzymes, OTUB1 and UCHL1. Consistent with these results, OTUB1 has recently been harnessed for targeted protein stabilization with deubiquitination-targeting chimeras (DUBTACs)(Henning et al., 2022a; Kanner et al., 2020). There were also other proteins involved in the ubiquitin-proteasome pathway, such as the ubiquitin-dependent aspartyl protease DDI1 (Yip et al., 2020) and TANK, an adaptor protein promoting the activity of the USP10 deubiquitinase (Wang et al., 2015). Other prominent hits with known roles in promoting protein stability included multiple chaperones and co-chaperones (e.g., HSPA1A, DNAJB3, CDC37, PDCL3, SGTA), calmodulin family proteins (CALM3 and CALML3), SUMOylation enzymes (UBE2I and UBA5), and four of the six 14-3-3 proteins present in the ORFeome (Bennett and Strehler, 2008; Celen and Sahin, 2020; Dar et al., 2014; Rosenzweig et al., 2019). However, most hits were not directly related to protein stabilization. Rather, they were enriched in proteins involved in diverse metabolic processes, such as nucleotide metabolism (**Figure S2D** and **S2E**). For example, the ribose-phosphate pyrophosphokinase 2 (PRPS2) was a robust hit in both stabilization screens. It is possible that proximity-dependent stabilizers can act both catalytically by removing ubiquitin (such as DUBs) or biophysically by increasing the thermodynamic stability of the EGFP-ABI1 fusion (e.g., PRPS2). Consistent with this possibility, stabilization screen hits were themselves significantly more stable than the rest of the ORFeome, as measured by protein half-life (Schwanhäusser et al., 2011) or by steady-state stability (Yen et al., 2008)(**Figures S2F-G**). In contrast, the half-life of degradation screen hits was not significantly lower than that of non-hits, and the difference in steady-state stability was modest (**Figures S2F-G**).

### E3 ligases have strikingly different activities in proximity-induced degradation

To comprehensively characterize the ability of E3 ligases to degrade targets in a proximity-dependent manner, we individually assessed the effect of 290 human E3 ligases fused to vhhGFP on the EGFP-ABI1 reporter stability in an arrayed screen format (**Figure 2**). Experiments were done in two independent replicates (**Figure S3A**). In line with the pooled ORFeome screen, not all E3 ligases efficiently promoted degradation of the reporter: 53 (18%) of the individually tested E3 ligase clones significantly decreased the EGFP/TagBFP ratio compared to the negative control Renilla luciferase (RLuc) fused to vhhGFP (**Figure 2** and **Table S1**). Importantly, the predicted E3 hits from the ORFeome screens were significantly better at degrading the reporter than non-hits (**Figure S3B**).

**Figure 2.**
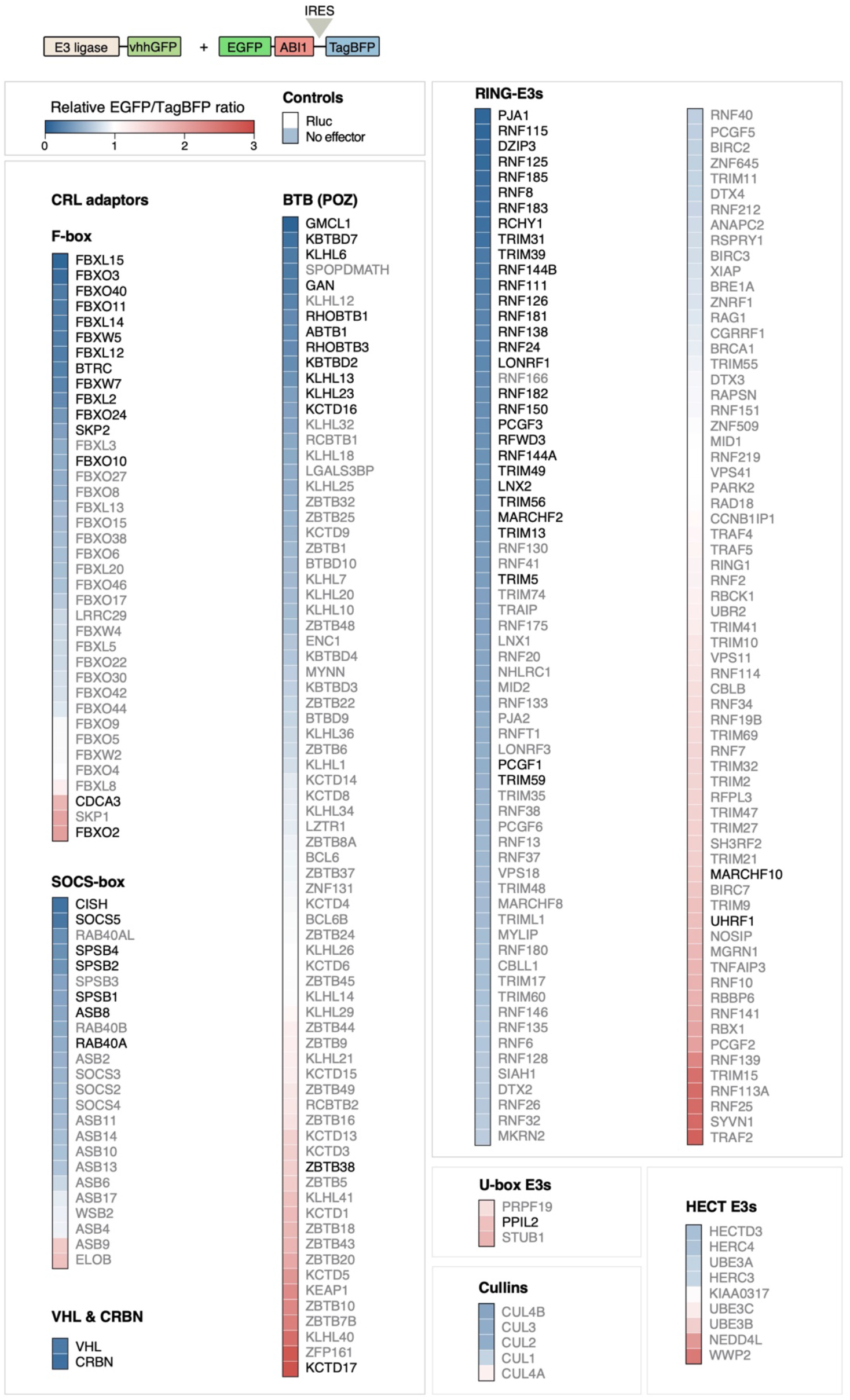
Characterization of human E3 ligases in proximity-dependent degradation assay. Top, 290 human E3 ligases fused to vhhGFP were transfected into 293T reporter cells expressing the EGFP-ABI1 reporter construct. Bottom, heat map of EGFP-ABI1 stability after E3-vhhGFP transfection. EGFP/TagBFP ratio was normalized to reporter cells transfected with Renilla luciferase (RLuc) fused to vhhGFP. E3 ligases that significantly decrease or increase the EGFP/TagBFP ratio compared to RLuc-vhhGFP are indicated in black. Statistical significance was calculated with an unpaired t-test and corrected for multiple hypothesis with the false discovery rate approach.

There were no statistically significant differences between different E3 ligase families in their ability degrade the reporter (**Figure S3C**). Consistent with this, we identified potent degradation effectors across all E3 families except the smaller HECT (Homologous to E6AP C-Terminus) and U-box families (**Figure 2**). However, there were significant differences in the activity of even highly related E3 ligases, as best illustrated by members of the FBXL or Kelch-BACK-BTB domain families (**Figure 2, S3D** and **S3E**), indicating that sequence similarity is not a robust predictor of activity in proximity-induced degradation. Notably, among the most potent proteins in this setting was CRBN, which was not part of the pooled ORFeome library (**Figure 2**). Among the 133 RING E3s tested, 33 significantly reduced GFP intensity. For three RING E3s (TRIM31, RCHY1, and RNF166), we also assayed alternative isoforms lacking a full-length RING finger. In all cases, loss of the RING finger led to loss of activity, indicating that these E3 ligases functioned through the expected E2-dependent ubiquitylation mechanism (**Figure S3F**). Together, these results establish that E3 ligases have vastly different potencies in inducing the degradation of an artificial substrate when brought to close proximity.

### E2 conjugating enzymes as proximity-dependent degradation effectors

One of the most potent degradation inducers that we identified in both ORFeome screens was the E2 conjugating enzyme UBE2B (also known as Rad6B)(**Figure 1E** and **1F**). UBE2B was a particularly interesting effector, because until recently, E2 enzymes had not been harnessed in targeted protein degradation (King et al., 2022). To further characterize UBE2B, we first tested if its catalytic activity is required for degradation. The active site of UBE2B contains Cys88 (**Figure 3A**), which is transiently conjugated to ubiquitin in a transthiolation reaction before ubiquitin transfer to the substrate or a HECT-type E3 ligase (Stewart et al., 2016). Mutating Cys88 to alanine (UBE2B^C88A^) completely abolished the activity without affecting protein stability, indicating that transthiolation is required for proximity-induced degradation by UBE2B (**Figures 3B** and **S4A**).

**Figure 3.**
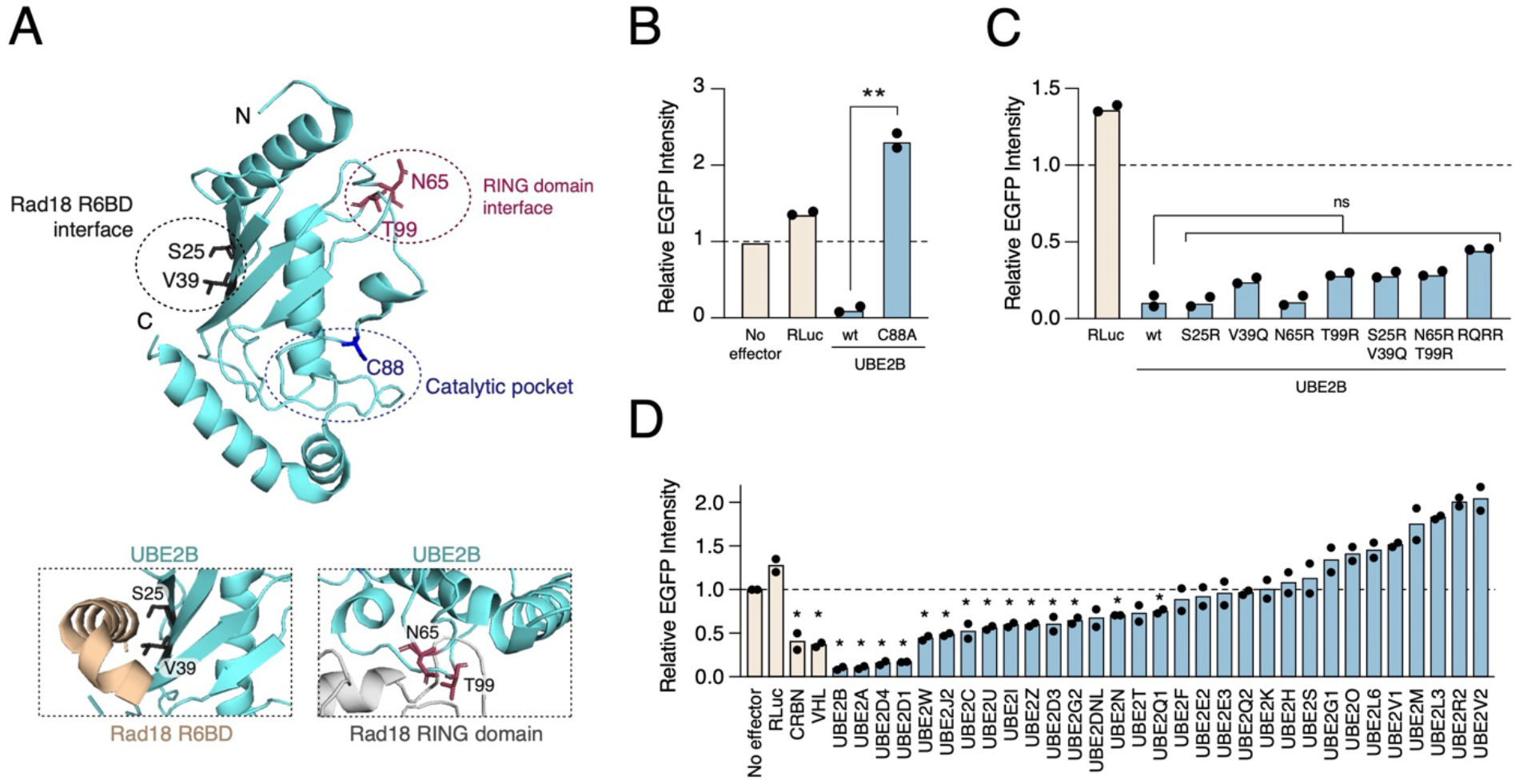
UBE2B is a potent proximity-dependent degrader. (**A**) Top, structure of UBE2B (cyan; PDB 2YB6). Residues mutated in the Rad18 R6BD interaction interface (black), the RING domain interaction interface (magenta), and in the catalytic pocket (blue) are indicated. Bottom, close-up views of the R6BD interface (Rad18 R6BD shown in light orange)(Hibbert et al., 2011) and a model of the RING interaction interface (Rad18 RING domain in grey). (**B**) Mutating the catalytic Cys88 to alanine abolishes the activity of UBE2B in the proximity degradation assay. The EGFP-ABI1 reporter cell line was transfected with indicated vhhGFP fusions and reporter stability was measured by flow cytometry. **(C)** Single or combined mutations in the E3 ligase interacting interfaces do not disrupt the activity of UBE2B in the degradation assay. RQRR: S25R/V39Q/N65R/T99R quadruple mutant. The experiment was performed as in panel (B). Statistical significance was calculated with an unpaired two-tailed t-test assuming equal variance and corrected for multiple hypotheses with false discovery rate correction. **(D)** Characterization of human E2 conjugating enzymes in proximity-dependent degradation. The EGFP-ABI1 reporter cell line was transfected with indicated vhhGFP fusions and reporter stability was measured by flow cytometry. Statistical significance was calculated with an unpaired two-tailed t-test assuming equal variance and corrected for multiple hypotheses with false discovery rate correction.

Because E2s function in tandem with E3 ligases in target ubiquitination, we next asked if UBE2B requires an E3 partner to promote proximity-dependent degradation. UBE2B is known to interact with the RING fingers of the E3 ligases RNF20 and RAD18 with the same interaction surface (Foglizzo et al., 2016; Hibbert et al., 2011; Kumar et al., 2015). RAD18 has also another domain, Rad6 binding domain (R6BD), which interacts with another surface on the opposite side of UBE2B (Kumar et al., 2015)(**Figure 3A**, bottom). We introduced disruptive single or double mutations in the interface that interacts with RAD18 and RNF20 RING fingers (N65R or T99R) or with RAD18 R6BD (S25R or V39Q). We also combined all four mutations into a single construct, UBE2B^RQRR^. All mutants were expressed at similar levels and were active in the degradation assay, suggesting that E3 binding is not required for the ability of UBE2B to degrade targets in a proximity-dependent manner (**Figures 3C** and **S4B**). This is consistent with a previous study showing that UBE2B can ubiquitinate a substrate protein *in vitro* in the absence of an E3, as long as the substrate is in close proximity (David et al., 2010).

To test if E2 conjugating enzymes are generally potent degraders, we measured the activity of 31 of the 38 human E2s in the degradation assay with EGFP-ABI1. Of these, 14 significantly reduced EGFP fluorescence (**Figure 3D** and **Table S1**). UBE2B was the most potent inducer of degradation together with its highly similar paralog UBE2A (**Figure 3D**). In addition, two related E2s UBE2D1 and UBE2D4 were also highly active degradation inducers, consistent with the recent discovery of a UBE2D-dependent molecular glue targeting NF-kB (King et al., 2022). Thus, despite the highly conserved E2 fold, there are clear differences in the functionality of E2s in mediating proximity-dependent degradation. We conclude that UBE2B, UBE2A, and UBE2D have an intrinsic potential as degradation effectors and surmise they could be harnessed for targeted protein degradation.

### Proximity-dependent degradation *in trans* via degron motifs

We identified several uncharacterized proteins as potent degraders in the screen. One of these was PRR20A, an uncharacterized proline-rich protein. It caught our attention because it does not contain any globular domains, as predicted by AlphaFold2 (Jumper et al., 2021). We therefore hypothesized that it might function indirectly by recruiting an E3 ligase or another factor through a degron sequence. This hypothesis was bolstered by the fact that another vhhGFP screen hit was EID1, an unstructured protein that contains a degron that binds the E3 ligase adaptor FBXO21 (Watanabe et al., 2015; Zhang et al., 2015).

Degrons can induce degradation of stable proteins when directly fused to them (i.e., *in cis*). We therefore fused full length PRR20A and EID1 or their fragments to EGFP in the vector that also expressed DsRed after the P2A ribosomal skipping motif. In this context, fluorescence ratio between the EGFP fusion and DsRed control acts as a stability reporter (**Figure 4A**, top). To compare degradation *in cis* and *in trans*, we also assayed the same constructs with the vhhGFP system (where EID1 and PRR20A are brought in proximity to EGFP intermolecularly). As expected, fusion of EGFP to the C-terminal degron fragment of EID1 led to low EGFP fluorescence, whereas the N-terminal fragment of EID1 fused to EGFP was stable (**Figure 4A**). The results were highly similar with the vhhGFP degradation assay, indicating that the EID1 degron can act both *in cis* and *in trans* (**Figure 4A**).

**Figure 4.**
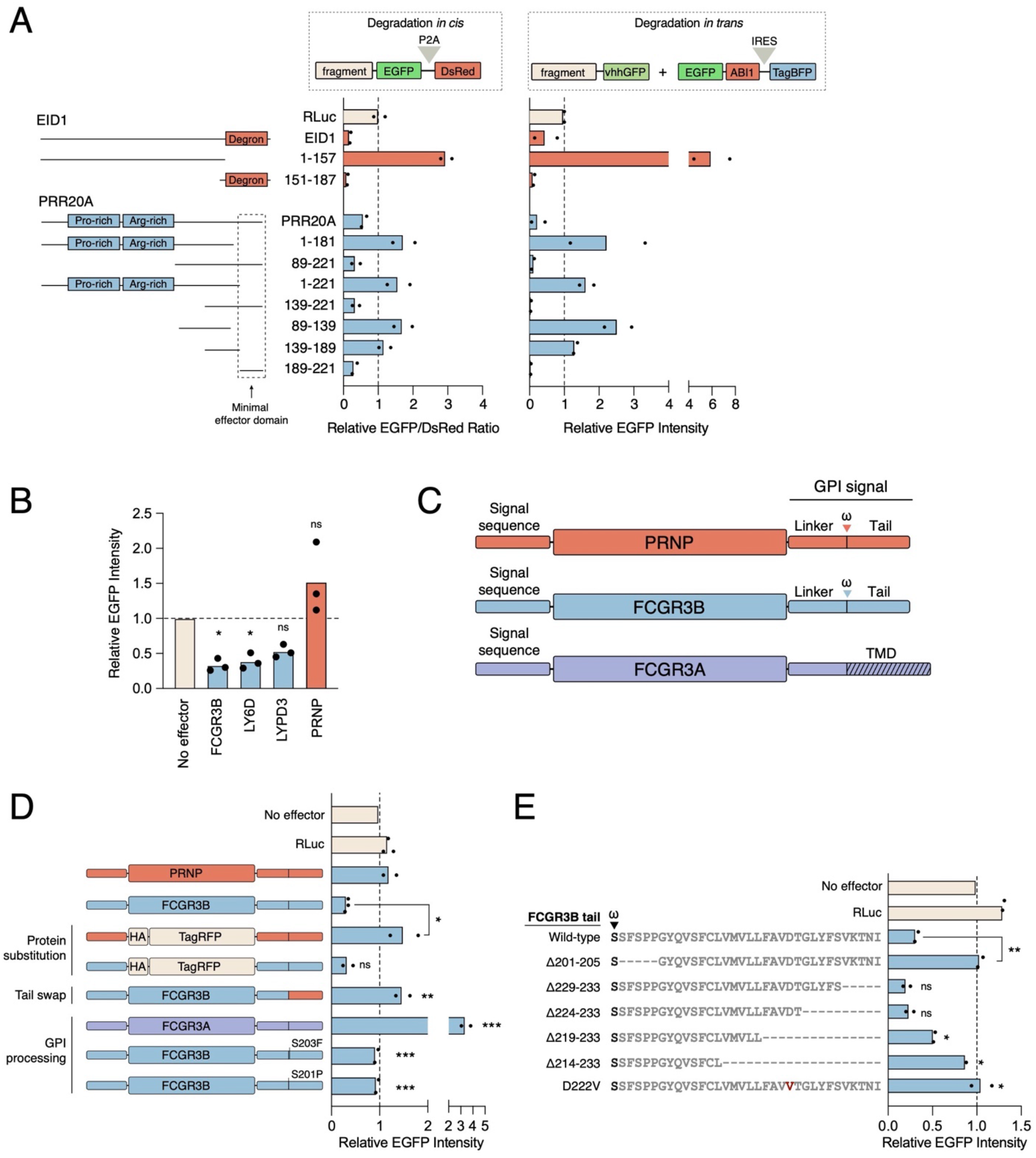
Degrons can function *in trans* for proximity-dependent degradation. (**A**) Full-length EID1 and PRR20A or their fragments were either directly fused to EGFP (left) to assess degradation *in cis* or to vhhGFP (right) for degradation *in trans*. EGFP fusions were transfected into 293T cells and protein stability measured as EGFP/DsRed ratio. VhhGFP fusions were transfected into the EGFP-ABI1 293T cell line. Fluorescence was measured with flow cytometry and normalized to Renilla luciferase (RLuc) fused to EGFP (left) or to RLuc-vhhGFP (right). (**B**) GPI-anchored proteins can mediate proximity-induced degradation. Indicated GPI-anchored proteins were transfected into the EGFP-ABI1 reporter cell line and EGFP levels were measured by flow cytometry. Fluorescence was normalized to reporter cells transfected with unrelated DNA (no effector). **(C)** Domain structures of FCGR3B, PRNP, and FCGR3A. ω is the residue to which the GPI moiety is attached. FCGR3A is not GPI anchored but contains a C-terminal transmembrane domain. (**D**) The EGFP-ABI1 293T reporter cell line was transfected with indicated vhhGFP fusion constructs. EGFP fluorescence was measured by flow cytometry and normalized to reporter cells transfected with unrelated DNA (no effector). (**E**) C-terminal deletion constructs of FCGR3B were assayed as vhhGFP fusions in the EGFP-ABI1 reporter cells as in (**D**). Statistical significance was calculated with one-way ANOVA with Dunnett’s multiple comparison test (*, p < 0.05; **, p < 0.01; ***, p < 0.001).

Full-length PRR20A fused directly to EGFP was also unstable. Using multiple overlapping fragments of PRR20A, we identified the C-terminal residues 189-221 as the region conferring low stability to EGFP fusions (**Figure 4A**, left panel). Similar to the EID1 degron, PRR20A^189-221^ could also degrade EGFP-ABI1 when directly fused to vhhGFP, indicating that the same region harbors both *in cis* and *in trans* degradation activity (**Figure 4A**, right panel). Thus, PRR20A likely acts in an indirect manner by recruiting a yet-unknown E3 ligase or another effector that degrades the target protein. More generally, these results indicate that our induced proximity screens not only identified E3 or E3-like degradation effectors but also uncovered trans-degrons that function indirectly on the target.

### Hydrophobic tails of GPI-anchored proteins can act as degrons

One highly enriched group of proteins in the degradation screens was GPI-anchored proteins, such as FCGR3B, LY6D, and LYPD3 (**Figures 1E-1H**). When tested individually as vhhGFP fusions, FCGR3B and LY6D degraded the EGFP-ABI1 reporter (**Figure 4B**). In contrast, another GPI-anchored protein, PRNP (prion protein) was neither a screen hit nor degraded the EGFP-ABI1 reporter when re-tested (**Figure 4B**). To identify the molecular determinants leading to this difference, we focused on FCGR3B (Fcγ receptor IIIb, also known as CD16b) and PRNP. Both proteins have a signal peptide that routes the protein to the secretory pathway, followed by a folded domain and a GPI anchor signal, which consists of a hydrophilic linker, cleavage site (ω), and a hydrophobic tail (Kinoshita, 2020)(**Figure 4C**). The hydrophobic tail is embedded in the membrane such that the vhhGFP moiety would be projected into the cytoplasm.

We first replaced the folded domains of PRNP and FCGR3B with TagRFP, while keeping their signal sequence and GPI anchor signal intact. These constructs behaved identically to the native proteins. Thus, the folded domains of these proteins – and by extension, their endogenous biological functions – do not have a role in EGFP-ABI1 degradation (**Figure 4D**). However, when we swapped the C-terminal hydrophobic tail of FCGR3B to that of PRNP, the construct failed to degrade EGFP (**Figure 4D**). This result indicates that the C-terminal hydrophobic tail is responsible for the difference between the activity of PRNP and FCGR3B in this context.

We next compared FCGR3B to its paralog FCGR3A, which almost identical in sequence but tethered to the plasma membrane through a transmembrane domain instead of a GPI anchor (Lanier et al., 1991)(**Figure 4C**). In contrast to FCGR3B, FCGR3A-vhhGFP fusion did not induce EGFP-ABI1 reporter degradation (**Figure 4D**). Notably, a single residue (Ser203 in FCGR3B vs Phe203 in FCGR3A) determines the mode of membrane attachment of these proteins (Lanier et al., 1991). Switching the residue in FCGR3B to that of FCGR3A (FCGR3B^S203F^) produced a protein that failed to degrade the EGFP-ABI1 reporter (**Figure 4D**), suggesting that GPI anchoring is involved in FCGR3B-induced degradation. Processing of the anchor includes proteolytic cleavage of the C-terminal hydrophobic tail prior to the conjugation of the GPI moiety (Kinoshita, 2020). Replacing the small ω+1 residue with a proline (FCGR3B^S201P^), which prevents cleavage, similarly rendered FCGR3B inactive in the assay (**Figure 4D**). Thus, the activity of FCGR3B-vhhGFP fusion in the degradation assay requires signal anchor cleavage, leaving the C-terminal tail fused to vhhGFP.

We hypothesized that the cleaved C-terminal tail might contain a degron sequence that promotes degradation of the cytoplasmic reporter. Therefore, we further defined the degradation-conferring region in FCGR3B with deletion constructs and alanine scanning. Consistent with the point mutants, deletion of five amino acids immediately following the ω site (Ser201-Pro205) abolished the activity of FCGR3B (**Figure 4E**). The C terminus of the hydrophobic tail was more tolerant to deletions, as removal of the last 10 amino acids did not notably affect the activity. However, removing 15 aa from the C terminus (or replacing amino acids Val221-Leu225 with alanines (**Figures 4E** and **S5**) significantly attenuated the ability of FCGR3B to degrade the reporter, suggesting that the center residues of the hydrophobic tail are functionally important. We noticed that this region contains a single negatively charged aspartate, which is a highly disfavored amino acid in single-pass transmembrane helices (Baker et al., 2017). Remarkably, changing this residue to a hydrophobic one (FCGR3B^D222V^) abolished the activity of the vhhGFP fusion (**Figure 4E**). These results suggest that mere hydrophobicity of the cleaved C-terminal tail of FCGR3B is not sufficient to drive degradation of a cytoplasmic protein in a proximity-dependent manner. Rather, the C-terminal tail likely contains a degron-like sequence that recruits a downstream effector for this function.

### Characterization of proximity-dependent stabilizers

We then turned our attention to proximity-dependent stabilizers. Like for E3 ligases, we first complemented the unbiased screen by individually assaying a panel of human deubiquitinases. However, because the original EGFP-ABI1 reporter is relatively stable, we designed a destabilized GFP reporter by fusing it to an unstable missense variant of glycine N-methyltransferase, GNMT^H176N^ (Luka et al., 2007; Sahni et al., 2015), followed by the P2A motif and DsRed (**Figure 5A**). We then individually co-transfected DUB-vhhGFP fusions with the GNMT^H176N^ reporter and assessed their effect on its stability (**Figure 5B**). Of the 99 DUBs encoded in the human genome (Clague et al., 2019), we tested 47, including ubiquitin-specific proteases (USPs), ubiquitin C-terminal hydrolases (UCHs), ovarian tumor proteases (OTUs), Machado-Josephin domain proteases (MJDs), zinc-dependent metalloenzymes (JAMMs) and SUMO proteases, as they encompass the majority of DUB superfamilies. Nine DUBs could significantly stabilize the reporter compared to the RLuc-vhhGFP control (**Figure 5B** and **Table S1**). These included the ubiquitin C-terminal hydrolases USP13, USP14, USP38, and USP39 as well as OTU proteases OTUB1 and OTUD6B. However, most DUBs had no effect on the unstable EGFP reporter. Thus, just like E3 ligases, DUBs differ greatly in their ability to regulate the stability of non-physiological substrates through induced proximity. This is likely a reflection of their intrinsic specificity towards distinct ubiquitin chains and topologies (Clague et al., 2019).

**Figure 5.**
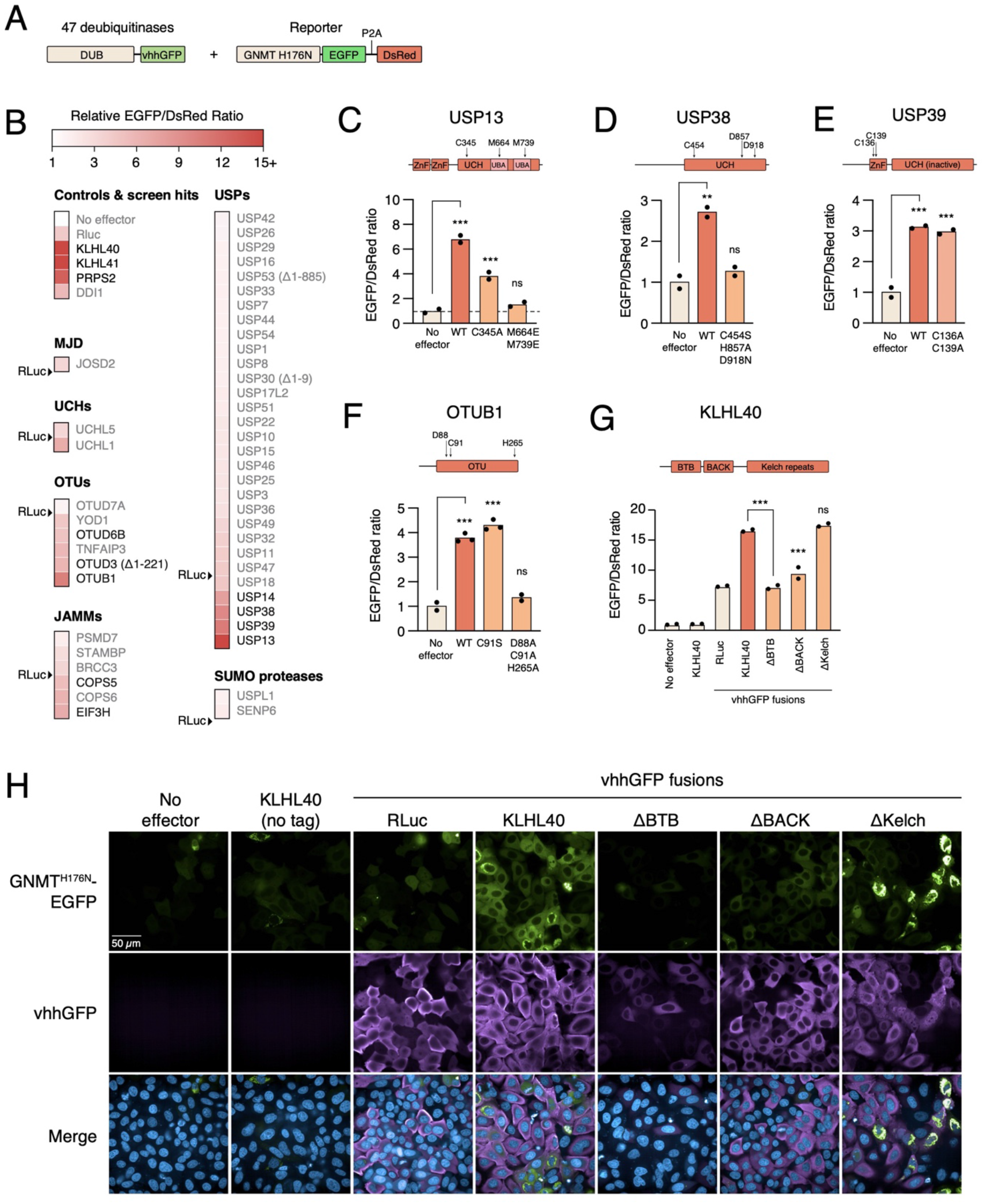
Deubiquitinases and KLHL40 as proximity-dependent stabilizers. **(A)** 47 human deubiquitinases fused to vhhGFP were co-transfected into 293T reporter cells with GNMT^H176N^-EGFP to assess their activity in proximity-dependent stabilization. **(B)**, Heat map of DUB activities in the stabilization assay. EGFP/DsRed ratio was normalized to cells transfected with an unrelated construct (no effector). Constructs that significantly increase the ratio are indicated in black. Statistical significance was calculated with an unpaired t-test and corrected for multiple hypothesis with false discovery rate correction. **(C-F)** Indicated mutants of USP13 (C), USP38 (D), USP39 (E), OTUB1 (F), and KLHL40 (G) were fused to vhhGFP and tested in the stabilization assay with GNMT^H176N^-EGFP. For KLHL40, an untagged construct without vhhGFP was also used as a control. Statistical significance was calculated with one-way ANOVA with Dunnett’s multiple comparison correction. *, p < 0.05; **, p < 0.01; ***, p < 0.001). (**H**) Microscopy images of HeLa cells transfected with GNMT^H176N^-EGFP and the indicated effector constructs. Cells were stained for the effector (vhhGFP fusion) with an anti-vhh antibody.

We then tested if catalytic activity was required for the ability of DUBs to stabilize the reporter. We mutated the catalytic residues or other functional residues of USP13, USP38, USP39, and OTUB1 and assessed the activity of the constructs in the stabilization assay (**Figures 5C-F** and **S6A**). Inactivating the catalytic cysteine of (USP13^C345A^) decreased the stabilizing effect of USP13, whereas disrupting the two ubiquitin-binding domains of USP13 (USP13^M664E/M739E^) (Zhang et al., 2011) abolished its activity on GNMT^H176N^, although both mutants were expressed at similar levels as the wild-type construct (**Figures 5C** and **S6A**). A catalytically dead mutant of USP38 (USP38^C454S/H857A/D918N^)(Chen et al., 2018) was similarly inactive in the assay (**Figure 5D**). Thus, these two DUBs appear to function through a mechanism requiring catalytic activity and ubiquitin binding. In contrast, USP39 has been reported to be a pseudoenzyme with no deubiquitinase activity *in vitro* (van Leuken et al., 2008; Walden et al., 2018). Moreover, mutating its ubiquitin-binding domain (USP39^C136A/C139A^) had no effect on activity (**Figure 5E**), suggesting that USP39 functions in an alternative manner in this context. Similarly, we observed that catalytic activity was dispensable for the function of OTUB1 in this assay (**Figure 5F**). However, OTUB1 also has an additional non-catalytic role in the cell, as it can stoichiometrically interact with and inhibit the activity of E2 enzymes (Juang et al., 2012; Nakada et al., 2010). To test this possibility, we assayed a triple mutant construct (OTUB1^D88A/C91A/H265A^) that is deficient in E2 binding and inhibition (Nakada et al., 2010) and also predicted to be deficient in K48-linked ubiquitin chain binding (Juang et al., 2012; Wiener et al., 2012). This mutant could not stabilize GNMT^H176N^-EGFP (**Figure 5F**), strongly suggesting that in proximity-dependent protein stabilization, OTUB1 acts via its E2 inhibiting function or its ability to bind to K48-linked ubiquitin chains rather than through its deubiquitinase activity.

### KLHL40 as a potent non-canonical stabilization effector

While assaying E3 ligases and deubiquitinases, we noticed that KLHL40 was an exceptionally potent stabilizer of both EGFP-ABI1 and GNMT^H176N^ (**Figures 2A** and **5B**). It was also a hit in both vhhGFP and PYL1 stabilization screens (**Figures 1I** and **1J**). Interestingly, previous work has indicated a role for KLHL40 in protein stabilization. Loss of KLHL40 leads to a decrease in the levels of its interacting partners nebulin (NEB) and LMOD3 (Garg et al., 2014), and in turn ectopic expression of KLHL40 can stabilize these proteins (Garg et al., 2014). KLHL40 belongs to the BTB-BACK-Kelch domain family, whose members normally function as adaptors for CUL3 CRLs (Canning et al., 2013). The BTB domain often interacts with CUL3 through a common φ-X-E motif (where φ represents a hydrophobic amino acid)(Canning et al., 2013). However, in KLHL40 this motif is not conserved (**Figure S6B**), suggesting that it does not function as a part of a CRL complex but through an alternative mechanism.

To characterize the function of KLHL40 in protein stabilization in more detail, we designed several deletion constructs. Full-length KLHL40 fused to vhhGFP stabilized the GNMT^H176N^ mutant reporter very efficiently as assessed by flow cytometry or microscopy (**Figures 5G and 5H**). Expressing KLHL40 without the vhhGFP moiety had no effect on GNMT^H176N^ stability, indicating that the activity was dependent on induced proximity (**Figures 5G** and **5H**). However, constructs lacking the BTB or the BACK domain could not stabilize the reporter (**Figures 5G** and **5H**), indicating that these domains are required for the stabilizing activity of KLHL40. In contrast, removing the Kelch domain (KLHL40^ΔKelch^) had no effect on the overall stability of GNMT^H176N^. Interestingly, however, GNMT^H176N^-EGFP stabilized by full-length KLHL40-vhhGFP appeared mostly diffuse in the cytoplasm, whereas the same construct stabilized by KLHL40^ΔKelch^ formed distinct clusters in the cell, reminiscent of aggregation (**Figure 5H**). This suggests that the KLHL40 Kelch domain does not contribute to target stabilization but may contribute to target solubilization.

Together, our results provide strong support to the previous finding that the endogenous function of KLHL40 is in protein stabilization, not degradation. Furthermore, this activity can be retargeted towards non-physiological substrates through induced proximity.

### Characterizing the mode-of-action of proximity-dependent effectors

To understand the mechanisms by which the proximity-dependent effectors regulate protein stability, we assembled a panel of 49 effectors from the pooled and arrayed screens. We individually transfected them as vhhGFP fusions into the original EGFP-ABI1 cell line. Consistent with the screen, most degradation screen hits degraded the reporter, whereas all three included stabilizer effectors (KLHL40, DDI1 and PRPS2) increased the EGFP signal (**Figure 6A** and **Table S1**). We then assessed how small molecule inhibitors of diverse protein homeostasis pathways modulate the activity of these effectors. To this end, we used the translation elongation inhibitor cycloheximide, the proteasome inhibitor bortezomib, the NEDDylation inhibitor MLN4924 (Soucy et al., 2009), and the VCP/p97 inhibitor CB-5083 (Anderson et al., 2015).

**Figure 6.**
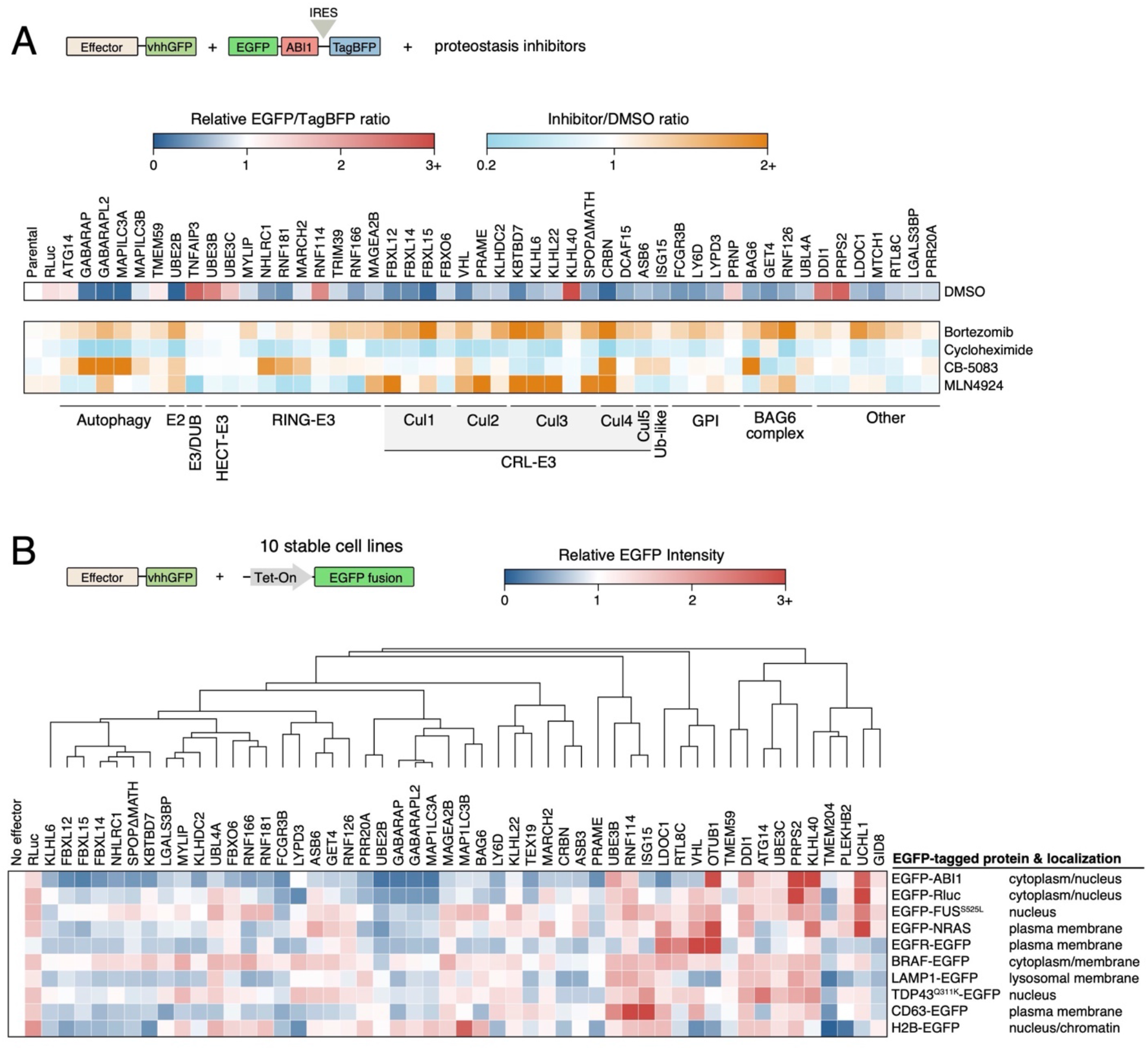
Characterizing the mode-of-action and target specificity of degradation and stabilization effectors. (**A**) Heatmap of relative EGFP/TagBFP ratios for a panel of 49 effectors selected from the pooled and arrayed screens. The EGFP-ABI1 reporter cell line was transfected with effectors fused to vhhGFP. Cells were treated with the indicated constructs and EGFP fluorescence was measured by flow cytometry. EGFP/TagBFP ratio was normalized to cells transfected with an unrelated DNA construct (no effector). For the inhibitor experiment, the heat map shows the relative change in EGFP/TagBFP ratio after inhibitor treatment. (**B**) 52 effectors were assessed for their activity against nine additional EGFP fusion proteins. Analysis was done as in panel (A). Effectors and targets were clustered using hierarchical clustering (one minus Pearson correlation and average linkage)

Inhibition of translation with cycloheximide treatment increased the degradative effect of almost all effectors, consistent with the global effect of the compound (**Figure 6A** and **Table S1**). Similarly, treatment of cells with bortezomib inhibited most degradation effectors, indicating that they were dependent on proteasome function (**Figure 6A**). In contrast, MLN4924, which specifically interferes with the function of CRLs, inhibited several CRL adaptors but had little effect on other proteins (**Figure 6A**). Interestingly, not all CRLs were equally affected by MLN4924. For example, FBXL14 was less affected by MLN4924 than the related F-box proteins FBXL12 and FBXL15. This is consistent with a previous study showing that a significant fraction of FBXL14 is associated with CUL1 even after MLN4924 treatment (Reitsma et al., 2017). Our results suggest that this remaining fraction might still be able to productively ubiquitinate substrates.

The VCP/p97 inhibitor CB-5083 also interfered with the degradation activity of some but not other effectors (**Figure 6A**). While it did not generally affect the ability of E3 ligases to degrade the reporter, it did specifically inhibit the function of CRBN. This is consistent with a recent report showing that VCP/p97 is required for degradation of CRBN substrates and neosubstrates (Nguyen *et al.,* 2017). Interestingly, another CUL4 CRL adaptor DCAF15 was not affected by CB-5083, indicating that even adaptors of the same CRL complex can promote protein degradation through distinct pathways. Another effector affected by CB-5083 was BAG6, which is a component of the BAG6/BAT3 complex that regulates the quality control of tail-anchored membrane proteins (Ganji et al., 2018). In line with this, VCP inhibition has been shown to induce stabilization and aggregation of BAG6 clients (Ganji et al., 2018). Thus, proximity-dependent degraders function mainly through their native mechanisms even when recruited to neosubstrates.

### Target specificity of stability effectors

Because our experiments had, until now, used only two reporters (EGFP-ABI1 and GNMT^H176N^-EGFP), we next asked if the identified effectors worked efficiently with multiple targets or if they were specific to the original reporter construct. To this end, we generated 9 stable, doxycycline-inducible 293T cell lines expressing diverse EGFP-tagged proteins localized to distinct cellular compartments and assayed the activity of 52 effectors as vhhGFP fusions in two biological replicates.

The effectors showed strikingly different patterns, with some being highly potent across multiple targets while others affected only a limited set of targets (**Figure 6B** and **Table S1**). For example, the E3 ligases FBXL12, FBXL14, FBXL15, and PRAME degraded almost all targets, whereas others such as ASB6, FBXO6, and RNF126 were much more selective. Other highly potent degradation inducers were the E2 conjugating enzyme UBE2B and the autophagy regulators MAP1LC3A, GABARAP, and GABARAPL2. Interestingly, Cereblon and VHL showed strikingly different effects. CRBN was a potent degrader of many targets (with the exception of FUS^P525L^ and NRAS), whereas VHL had a very limited target spectrum (**Figure 6B**). We observed similar target-specific differences for proximity-dependent stabilizers. For example, UCHL1 and OTUB1 deubiquitinases were much more selective in their target spectrum than KLHL40 or DDI1, which stabilized nearly all targets (**Figure 6B**).

Looking at the effector/target matrix in more detail, we noticed that several effectors preferred distinct classes of targets. For example, the BAG6/BAT3 complex members BAG6 and UBL4A preferentially degraded membrane proteins, in line with the role of the complex in membrane protein quality control. Similarly, ATG14 specifically degraded targets localized to the plasma membrane (EGFR, NRAS, CD63). ATG14 is a subunit of the autophagy-specific PI3 kinase complex (Dikic and Elazar, 2018; Matsunaga et al., 2010), and it promotes phagophore nucleation, tethering of autophagic membranes, and fusion of autophagosomes to endolysosomes (Diao et al., 2015). Thus, its activity towards membrane-associated proteins is in line with its known function in autophagy. As membrane proteins have been challenging targets for PROTACs that exploit cytoplasmic E3 ligases like VHL or CRBN, these proteins could represent more optimal effectors for targeted protein degradation of membrane associated proteins (Békés et al., 2022).

Looking at the specificity from the target side, each EGFP-tagged protein identified a unique complement of effectors that worked. This included constructs that were localized even to the same compartments, such as the EGFP-ABI1 and EGFP-Renilla luciferase fusions. Targets also differed in their sensitivity to proximity-induced modulation of stability. For example, 29 different effectors reduced EGFR-EGFP levels more than 25%, whereas only three (UBE2B, TMEM204, GID8) did so for BRAF-EGFP (**Figure 6B**). These results are in line with the observations that some targets appear to be intrinsically more susceptible to targeted degradation (Donovan et al., 2020). More broadly, these results indicate that both effectors and targets have clear pairing preferences, highlighting the complex interplay in proximity-dependent modulation of protein stability.

### Effector sensitivity to tag location

Development of potent heterobifunctional degraders is often complicated by the sensitivity of these molecules to the topology of the E3-target interaction. In many cases, minor changes in linker chemistry can modulate or even completely abolish the activity of the molecule in an unpredictable manner (Békés et al., 2022; Burslem and Crews, 2020; Donovan et al., 2020; Posternak et al., 2020). To test if the effectors identified in our screen are limited to specific geometries, we tagged 38 effectors with either N-terminal or C-terminal vhhGFP and assayed their activity in the original reporter cell line (**Figure S7**). While several effectors (such as SPOP, TRIM39, and LY6D) only worked as C-terminal fusions, others (such as UBE2B, FBXL12, and FBXL14) were equally potent regardless of the tag location (**Figure S7**). We suggest that effectors that are not sensitive to tag location would be ideal targets for the development of novel heterobifunctional molecules, as they may be more tolerant to different geometries.

### Recruiting effectors via diverse affinity tags

Until now, all our experiments were conducted with EGFP tagged proteins and either vhhGFP or PYL1 recruitment, leaving open the possibility that the screen had identified effectors that are dependent on these components for their activity. Indeed, a previous study described a covalent K-Ras PROTAC that can degrade K-Ras^G12C^ as an EGFP fusion but has no activity towards the endogenous protein (Zeng et al., 2020). Therefore, we tested if the effectors are functional also when they are brought to the target by alternative means. To do so, we first used an intracellular single domain antibody (iDab) that binds Ras family proteins with high affinity (Tanaka and Rabbitts, 2003). It has also been used for targeted degradation of K-Ras as a fusion to VHL or to the U-box of STUB1 E3 ligase (Bery et al., 2020). Due to the difficulty of detecting endogenous Ras proteins, we co-transfected 3xFLAG-V5 tagged K-Ras with the effectors fused to the binders As previously reported, STUB1^U-box^ fused to Ras iDab reduced K-Ras levels compared to an iDab targeting LMO2, an unrelated protein (Bery et al., 2020)(**Figure S8A**). Importantly, most degradation effectors similarly downregulated K-Ras protein, whereas KLHL40 stabilized it, consistent with the results with EGFP fusion proteins. Thus, most effectors are compatible with multiple different methods for induced proximity and not merely targeting the EGFP moiety.

Recently, a comprehensive study characterized the effect of 91 PROTACs on human protein kinases (Donovan et al., 2020), revealing a striking difference in the susceptibility of kinases to targeted protein degradation. While some kinases, such as CDK4 and WEE1, were degraded with many PROTACs, others like ARAF and IKBKE were not degraded by any compounds that engage VHL or CRBN. To test if some of our novel effectors would be able to degrade these targets, we tagged them with the 13-aa ALFA tag that is recognized by the NbALFA nanobody with high affinity (Götzke et al., 2019). Consistent with the chemical proteomics approach, we observed that VHL-NbALFA could not degrade either kinase in this assay (**Figure 7A** and **S8B**). (We could not express CRBN-NbALFA fusion at detectable levels.) In contrast, many hits from the ORFeome screen were much more efficient. For example, FBXL12, FBXL15, KLHDC2 and the GPI-anchored protein FCGR3B fused to NbALFA potently lowered the levels of ARAF, whereas KLHL40 increased the levels (**Figure 7A**). While IKBKE was generally more resistant to degradation than ARAF in this context, several proteins such as FBXL12, FBXL15, and FCGR3B reduced IKBKE levels, in contrast to VHL that had no effect (**Figure S8B**).

**Figure 7.**
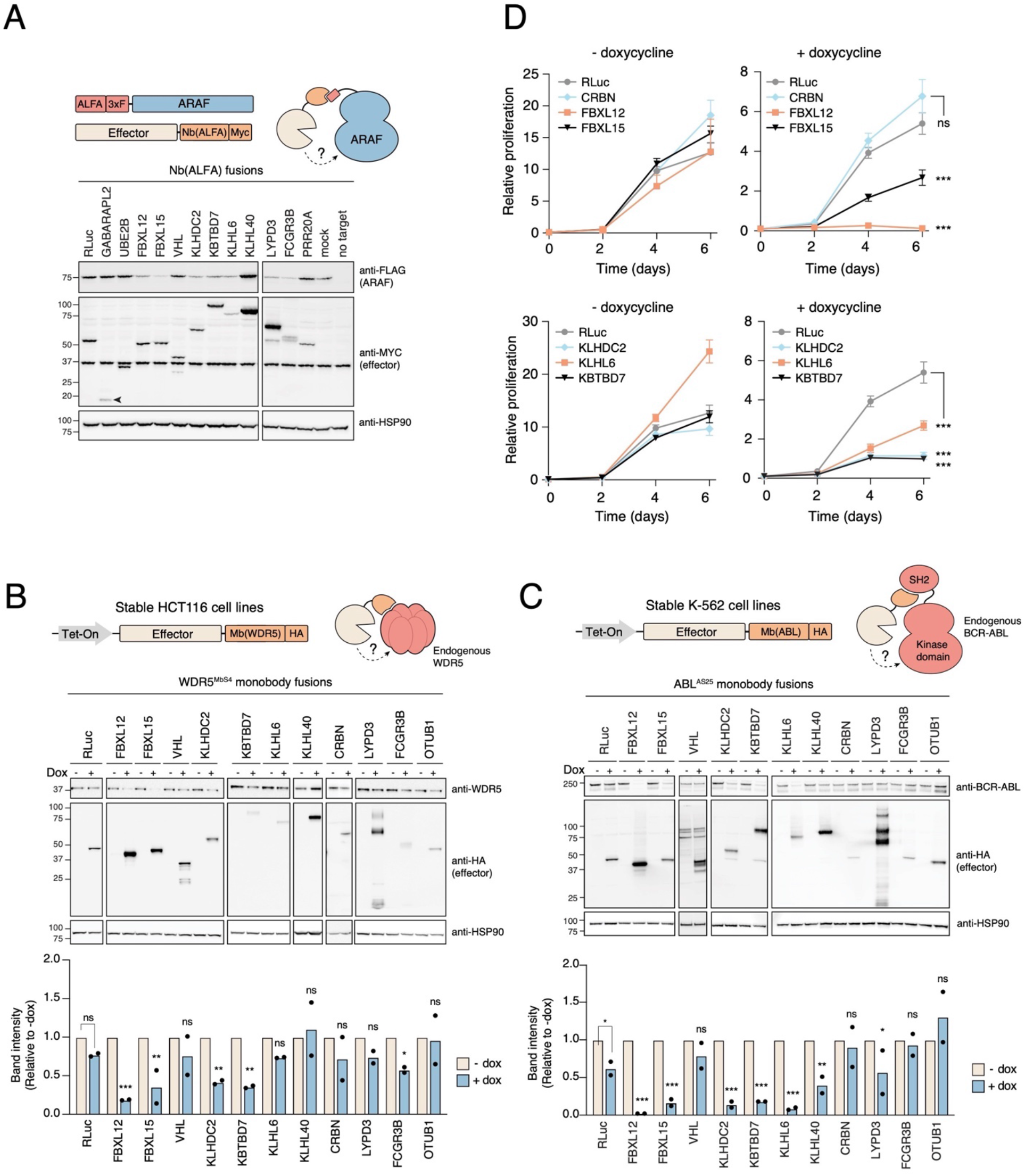
Benchmarking novel effectors with multiple recruitment strategies and therapeutically relevant targets. (**A**) Indicated effectors fused to Nb(ALFA)-Myc were co-transfected with ALFA-3xFLAG-ARAF into 293T cells, followed by western blotting for ARAF (anti-FLAG), effector (anti-Myc), and Hsp90. (**B**) Stable HCT116 cell lines expressing doxycycline-inducible effectors fused a WDR5-targeting monobody Mb(WDR5) were treated with doxycycline or left untreated. Top, Endogenous WDR5 levels and effector expression were assessed by western blotting. Bottom, Quantification of WDR5 levels after doxycycline induction. Statistical significance was calculated with an unpaired t-test with false discovery rate correction for multiple hypotheses. (**C**) As in (B), but effectors were fused to a monobody binding the SH2 domain of BCR-ABL and stably expressed in K562 cells. (**D**) K562 cell proliferation after induction of indicated effector fusion constructs with doxycycline. Note that the monobody itself has an effect on cell proliferation (compare top left graph scale to top right graph for RLuc-vhhGFP).

### Targeting endogenous proteins with novel effectors

Finally, we assessed the potency of the effectors in regulating the levels of endogenous proteins. To this end, we employed monobodies against the chromatin regulator WDR5 and the fusion oncogene BCR-ABL (Gupta et al., 2018; Wojcik et al., 2016). We cloned selected effectors into a doxycycline-inducible lentiviral vector with a C-terminal fusion to either monobody and generated stable HCT116 cells (for WDR5) and K562 cells (for BCR-ABL). We used VHL and CRBN as controls. While VHL could not degrade endogenous WDR5 and CRBN only had a modest effect, several novel effectors robustly degraded WDR5 in a doxycycline-dependent manner (**Figure 7B**). In particular, similar to the ALFA tag experiments, FBXL12 and FBXL15 were highly efficient in degrading endogenous WDR5. The results with BCR-ABL in K562 cells were similar: FBXL12, FBXL15, KBTBD7, KLHDC2, and KLHL6 degraded BCR-ABL highly efficiently, whereas CRBN and VHL had no significant effect (**Figure 7C**).

Because BCR-ABL is essential for the growth of K562 cells, we then assessed whether different effector fusions would have distinct effects on cell proliferation. Although the monobody alone inhibits the function of BCR-ABL (Wojcik et al., 2016), it only partially inhibited the proliferation of K562 cells in our system when fused to Renilla luciferase (**Figure 7D**). However, when the monobody was fused to FBXL12, FBXL15, KLHLDC2, KLHL6, or KBTBD7, the cells proliferated significantly slower than in the control condition or they ceased proliferating altogether (**Figure 7D**). In contrast, CRBN did not affect proliferation more than Renilla luciferase (**Figure 7D**). Consistent with these results, we observed a significant correlation between the ability of the effectors to degrade BCR-ABL and their effect on cell proliferation (**Figure S8C**). Thus, effectors identified in our unbiased ORFeome screen are significantly better at degrading and inhibiting the function of the hallmark oncogenic fusion of chronic myeloid leukemia, BCR-ABL.

## DISCUSSION

Our results suggest that the human proteome contains a significant cache of proteins that can regulate protein stability *in trans* in a proximity-dependent manner. This catalogue of functional effectors could be prioritized for ligand discovery and subsequently harnessed for targeted protein degradation or stabilization. At the same time, the results provide insights into the functional differences within central protein families regulating protein homeostasis.

### Inherent differences in E3 ligase and DUB efficacy

Our work revealed vast differences in the ability of E3 ligases to degrade and deubiquitinases to stabilize non-physiological target proteins. Indeed, only a relatively small fraction of E3s and DUBs could modulate the levels of our model substrates in a proximity-dependent manner. Several mechanisms could explain these differences. The first is the inherent specificity and activity of these proteins. Both E3 ligases and DUBs have a preference for specific types of ubiquitin linkages, and only some linkage types are associated with degradation by the proteasome (Clague et al., 2019; Swatek and Komander, 2016). Along the same lines, some E3s and DUBs prefer conjugating or cleaving monoubiquitin over polyubiquitin chains, which are predominantly (although not exclusively) responsible for degradation (Braten et al., 2016; Swatek and Komander, 2016). Moreover, some E3 ligases can ubiquitinate zones of lysines in a target protein, whereas others are specific for lysines in a particular context (Cowan and Ciulli, 2022; Mattiroli and Sixma, 2014). There are also vast differences in the catalytic activity of E3s and DUBs *in vitro* and *in vivo*, where the activity is often regulated by conformational changes, inhibitory domains, and post-translational modifications (Chen et al., 2017; Clague et al., 2019; Trempe et al., 2013). Several E3s and DUBs are actually pseudoenzymes that lack any activity (Clague et al., 2019; Walden et al., 2018). It is likely that this diversity in intrinsic specificity and activity is directly reflected in the activity of E3s and DUBs in our assay.

The second level of differentiation likely comes from the geometry of the effector-target complex. Structural studies with PROTACs and molecular glues have established that contacts between the E3 and the target are critical for stabilizing the ternary complex and for compound activity (Casement et al., 2021; Fischer et al., 2014; Gadd et al., 2017). Indeed, even minor changes in linker chemistry can have profound effects on PROTAC activity, and converting promiscuous inhibitors into degraders often dramatically limits their target specificity (Bondeson et al., 2018; Nguyen et al., 2021; Posternak et al., 2020). In a similar manner, we observed that some effectors were highly dependent on the location of the proximity tag (C vs N terminus) or the target identity or location. At the same time, other effectors were highly promiscuous across multiple targets and insensitive to tag location, indicating that their activity is less sensitive to extrinsic factors.

Developing effective PROTACs or molecular glues is a resource intensive pursuit, and thus the effector-target pairing must be carefully considered. Although different from the native context, our approach allows evaluating the potential activity of effectors and their compatibility with a single or multiple targets. We suggest that highly potent and promiscuous effectors identified in our screen (such as FBXL12, FBXL15, KLHDC2, or KBTBD7) may provide a more robust platform for the development of novel targeted protein degradation therapeutics, as these effectors are likely to be less sensitive to linker chemistries, target localization, and other parameters. On the other hand, picking more specific effectors that are expressed in a limited number of tissues (Jevtić et al., 2021) could alleviate potential off-target effects of next generation PROTACs and DUBTACs.

### *In trans* degradation by degrons

Our screen also uncovered degrons that were able to degrade a model substrate in a proximity-dependent manner. Typically, degrons induce degradation when they are part of a polypeptide chain (*in cis*) (Guharoy et al., 2016). However, in our case recruitment of the degron in PRR20A, a poorly characterized protein, led to collateral degradation of the proximal target when part of a different polypeptide chain (*in trans*). This is analogous to PROTACs that engage E3 ligases by mimicking their cognate degrons, such as the HIF1a degron for VHL or the IκBα degron for β-TRCP (Bondeson et al., 2015; Jevtić et al., 2021; Sakamoto et al., 2001). Identifying the E3 ligase or other effectors that interact with the PRR20A degron could point to novel effectors amenable to degron hijacking for targeted protein degradation. Moreover, just like screens of peptide libraries fused to EGFP have revealed *in cis* degrons (Koren et al., 2018; Timms et al., 2019), induced-proximity screens with similar libraries fused to vhhGFP or PYL1 could reveal many more short motifs that act *in trans*.

### UBE2B and E2 conjugating enzymes as proximity-dependent degraders

One of the most robust degraders we identified was the E2 conjugating enzyme UBE2B. As was the case for E3s and DUBs, E2s also showed striking differences in their activity in the degradation assay, indicating high functional diversity even within this highly conserved and compact protein family. The promiscuous activity of UBE2B towards diverse and unrelated targets indicates that it could be a powerful effector for targeted protein degradation applications. While E2s have not been considered easily druggable due to the lack of deep pockets, recent reports have demonstrated that they can be targeted at allosteric sites or by molecular glues that promote their interactions with other proteins (Hewitt et al., 2016; Huang et al., 2014; King et al., 2022; Morreale et al., 2017; St-Cyr et al., 2021). These results suggest that hijacking E2s for targeted protein degradation may not be as challenging as previously thought.

### Discovery of unexpected degradation and stabilization effectors

Many hits we identified in the unbiased screens did not fall into the expected categories in the ubiquitin-proteasome system or the protein homeostasis network at large. Indeed, we discovered several unexpected or even counterintuitive effectors. A notable class of proximity-dependent degraders consisted of GPI-anchored proteins (GPI-APs), which are cell surface proteins attached to the membrane via the glycosylphosphatidylinositol moiety. These proteins were not degraders due to their intrinsic activity, but rather due to the cleavage of their C-terminal signal peptide. This leaves a hydrophobic transmembrane domain attached to the N-terminus of the cytoplasmic proximity tag (vhhGFP or PYL1) used in our assay. Transmembrane domains can act as degrons when fused to a cytoplasmic protein (Mashahreh et al., 2022), and it is possible that the cleaved GPI signal peptide acts as a hydrophobic *trans*-degron in our assay setup. Previous work has shown that defects in GPI anchor biogenesis lead to the degradation of the misprocessed protein by multiple routes (Zavodszky and Hegde, 2019). Moreover, the cleaved GPI signal peptide appears to be degraded through the proteasome (Guizzunti and Zurzolo, 2014). It is therefore likely that proximity-dependent degradation of cytoplasmic proteins by GPI signal peptides occurs via the native quality control system of GPI-APs. Interestingly, recent cryo-EM structure of the GPI transamidase identified the E3 ligase RNF121 as an integral component of this multiprotein complex (Zhang et al., 2022), suggesting that it may be involved in this quality control step.

Another seemingly unexpected finding was that KLHL40, which belongs to the CUL3 E3 ligase adaptor family, acted as a potent stabilizer in a proximity-dependent manner. In contrast, most members of this family regulate protein ubiquitination and proteasome-dependent degradation (Canning et al., 2013; Stogios and Privé, 2004; Stogios et al., 2005). However, the native function of KLHL40 appears to be stabilization of its interacting partners LMOD3 and NEB by preventing their ubiquitination (Garg et al., 2014). These three proteins are specifically expressed in the skeletal muscle cells, where they are key regulators of thin filaments. Consistent with this, loss-of-function mutations in these genes cause nemaline myopathy (Lehtokari et al., 2014;

Ravenscroft et al., 2013; Yuen et al., 2014). While the exact mechanisms by which KLHL40 stabilizes its targets remain to be determined, our results raise the exciting possibility that KLHL40 could be hijacked for targeted protein stabilization in muscle tissues. This would be particularly relevant for the myriad of myopathies and muscular dystrophies, which are often caused by destabilizing mutations in large genes that are not amenable to gene replacement therapy (Dowling et al., 2021; Maani et al., 2021).

### Harnessing induced proximity beyond protein stability

The clinical success of molecular glues like lenalidomide and the promising early clinical results with PROTACs have amply demonstrated the therapeutic potential of targeted protein degradation (Békés et al., 2022; Deshaies, 2020). This has also spurred the development of other heterobifunctional molecules that promote target degradation through other mechanisms, such as autophagy (AUTACs), lysosomes (LYTACs), or nucleases (RIBOTACs)(Ahn et al., 2021; Banik et al., 2020; Dey and Jaffrey, 2019; Takahashi et al., 2019). But nature has demonstrated that the potential of induced proximity extends well beyond target degradation. Many plant hormones (such as abscisic acid) function by inducing specific protein-protein interactions, viral proteins hijack host proteins for their own advantage, and natural products such as rapamycin and brefeldin A act as molecular glues between two proteins to inhibit core cellular pathways (Gerry and Schreiber, 2020).

Exploiting the whole proteome and the entire complement of molecular pathways for induced proximity would offer an enormous opportunity for therapeutic intervention (Deshaies, 2020; Gerry and Schreiber, 2020; Modell et al., 2021). Indeed, proof-of-principle experiments have established that the activity of e.g. phosphatases and kinases can be re-targeted with heterobifunctional molecules (Modell et al., 2021; Siriwardena et al., 2020; Yamazoe et al., 2020). However, a major challenge is to identify which proteins in the proteome would have the desired effect on a given target, when induced to interact. Current hypothesis-driven approaches for induced proximity are strongly biased towards for known mechanisms and effectors instead of considering the entire proteome. As a case in point, it is doubtful that rapamycin would have been developed by rational design, as it brings together two unrelated proteins (mTOR and FKBP12) that have no functional connection in normal cells.

We suggest that our approach, which surveys almost the entire proteome for functional, proximity-dependent outcomes, can solve this “protein pair problem” and identify highly potent effectors for almost any therapeutically relevant target or process. In this study, we specifically screened for protein degraders and stabilizers, but the approach can be used to discover many other kinds of effectors. Indeed, we and others have used induced proximity screens to identify effectors of transcription (Alerasool et al., 2022; Stampfel et al., 2015; Tycko et al., 2020), RNA metabolism (Luo et al., 2020), and signal transduction (DeVit et al., 2005). Expanding the approach to therapeutically important targets such as oncogenes, tumor suppressors, and key immune regulators could provide many novel and unexpected protein pairs for targeting by molecular glues and heterobifunctional molecules. Together with recent advances in the discovery, development, and characterization of molecular glues (Domostegui et al., 2022; Mayor-Ruiz et al., 2020; St-Cyr et al., 2021) and covalent binders (Abbasov et al., 2021; Kuljanin et al., 2021; Parker et al., 2017; Spradlin et al., 2021; Vinogradova et al., 2020), induced-proximity proteomics could provide a springboard for the development of next-generation therapeutics.

## ACKNOWLEDGEMENTS

We would like to thank Zoheyr Imrit and Chris Mogg for their help with experiments and Daniel Durocher, Frank Sicheri, and the Taipale lab for their comments on the manuscript. This work was supported by the David Dime and Elisa Nuyten Catalyst Fund award for Mikko Taipale, the Mark Foundation for Cancer Research ASPIRE Award to Mikko Taipale (co-PIs: F. Sicheri and D. Durocher), the Charles H. Best Postdoctoral Fellowship to Dr. Juline Poirson, and the CIHR Fellowship Award to Dr. Hanna Cho.

## AUTHOR CONTRIBUTIONS

Conceptualization: J.P., A.D., N.A., L.M., M.T.; Investigation: J.P., A.D., H.C., M.L., N.A., J.L., L.M., M.T.; Formal Analysis: J.P., H.C., M.T.; Visualization: J.P., H.C., M.T.; Writing – Original Draft: J.P. and M.T.; Writing – Review & Editing: J.P., H.C., M.T.; Supervision: M.T.; Funding Acquisition: M.T.

## SUPPLEMENTARY FIGURES

**Figure S1.**
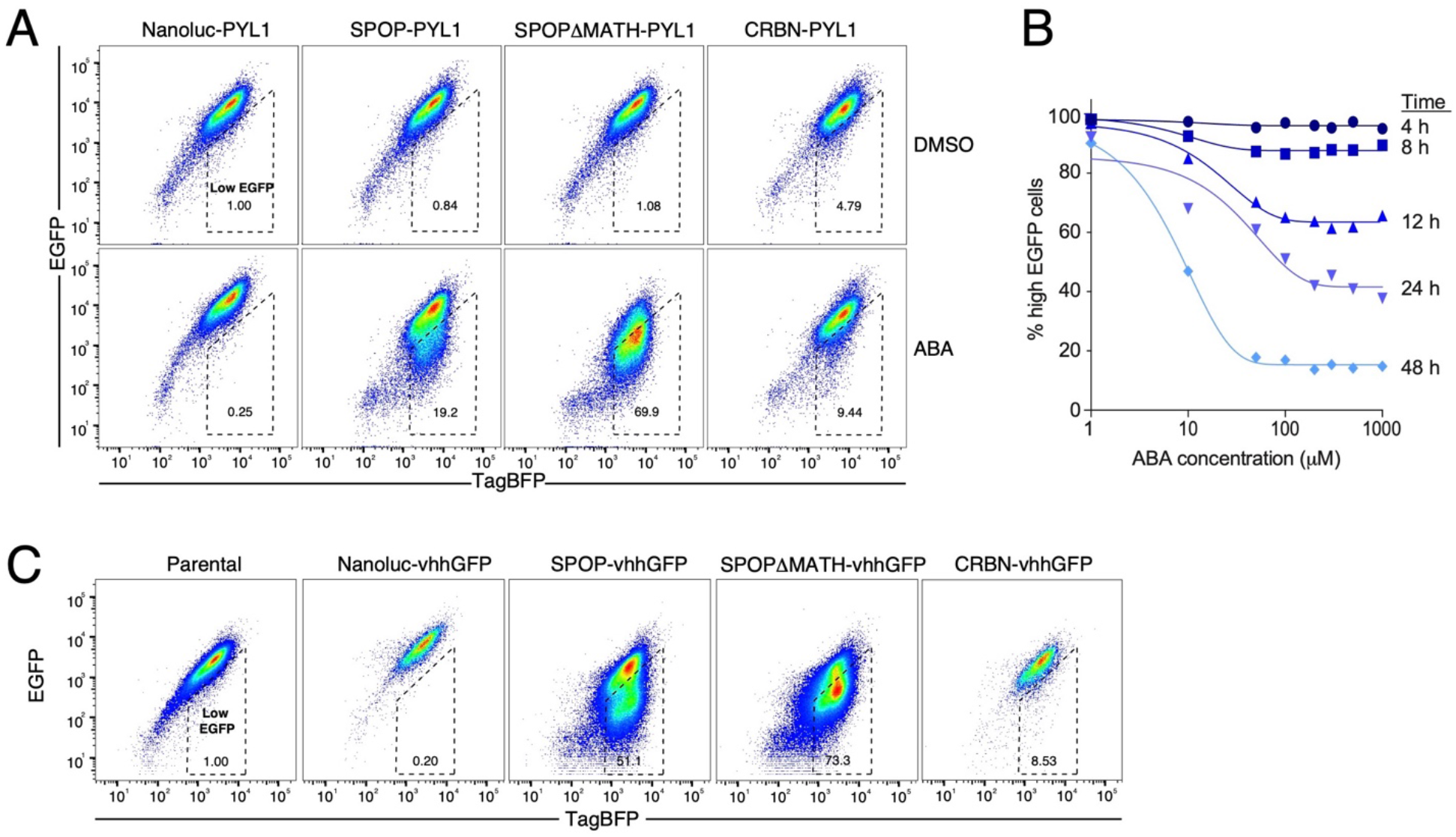
Establishing the induced proximity assay. Related to Figure 1. (**A**) Representative flow cytometry plots of the EGFP-ABI1-IRES-TagBFP dual-reporter 293T stable line transduced with C-terminal PYL1 fusions of Nanoluc, CRBN, full-length SPOP, or mutant SPOP lacking its native substrate-binding domain (residues 28-166, ΔMATH). Cells were then treated with either 0.5% DMSO or 100µM abscisic acid (ABA) for 24 hours. (**B**) Frequency of high EGFP population as a function of abscisic acid (ABA) concentration and treatment time in the EGFP-ABI1 293T reporter cell line transduced with the C-terminal SPOPΔMATH-PYL1 fusion. (**C**) Representative flow cytometry plots of the EGFP-ABI1-IRES-TagBFP dual-reporter 293T stable line transduced with C-terminal vhhGFP fusions of Nanoluc, CRBN, full-length SPOP, or SPOPΔMATH.

**Figure S2.**
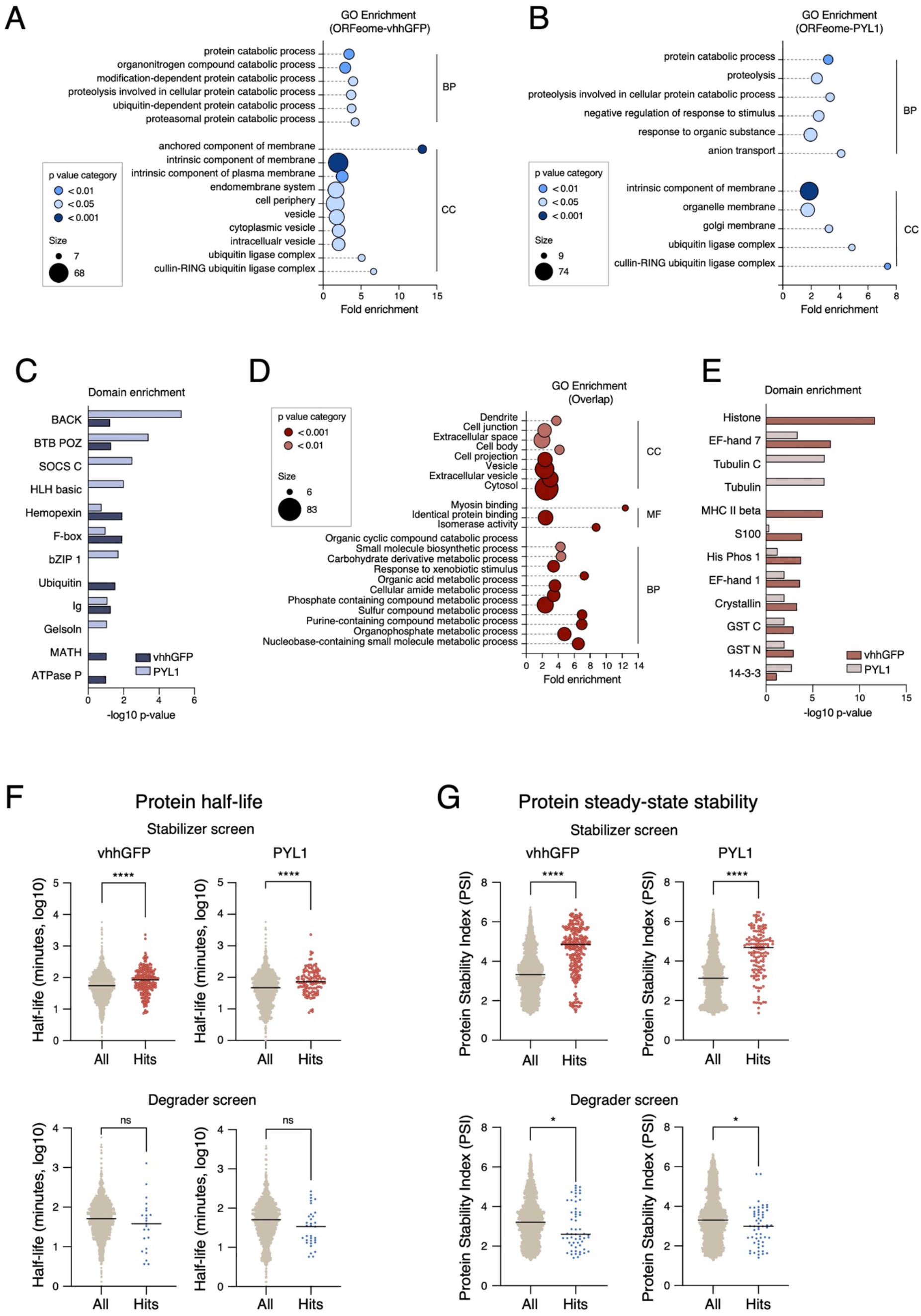
Features of effector proteins identified in pooled screens. Related to Figure 1. **(A-B)** Enrichment of Gene Ontology (GO) categories in the ORFeome-vhhGFP screen hits (A) and ORFeome-PYL1 screen hits (B). (**C**) Enrichment of PFAM and Interpro domains in the degradation screen hits. (**D**) Enriched GO categories in the overlapping hits between ORFeome-vhhGFP and ORFeome-PYL1 stabilization screens. (**E**) Enrichment of PFAM and Interpro domains in the stabilizer hits. (**F-G**) Comparison of protein half-lifes (Schwanhäusser et al., 2011)(F) and protein steady state levels (Yen et al., 2008) (G) between hits and all tested proteins. Statistical significance was calculated with a two-tailed t-test.

**Figure S3.**
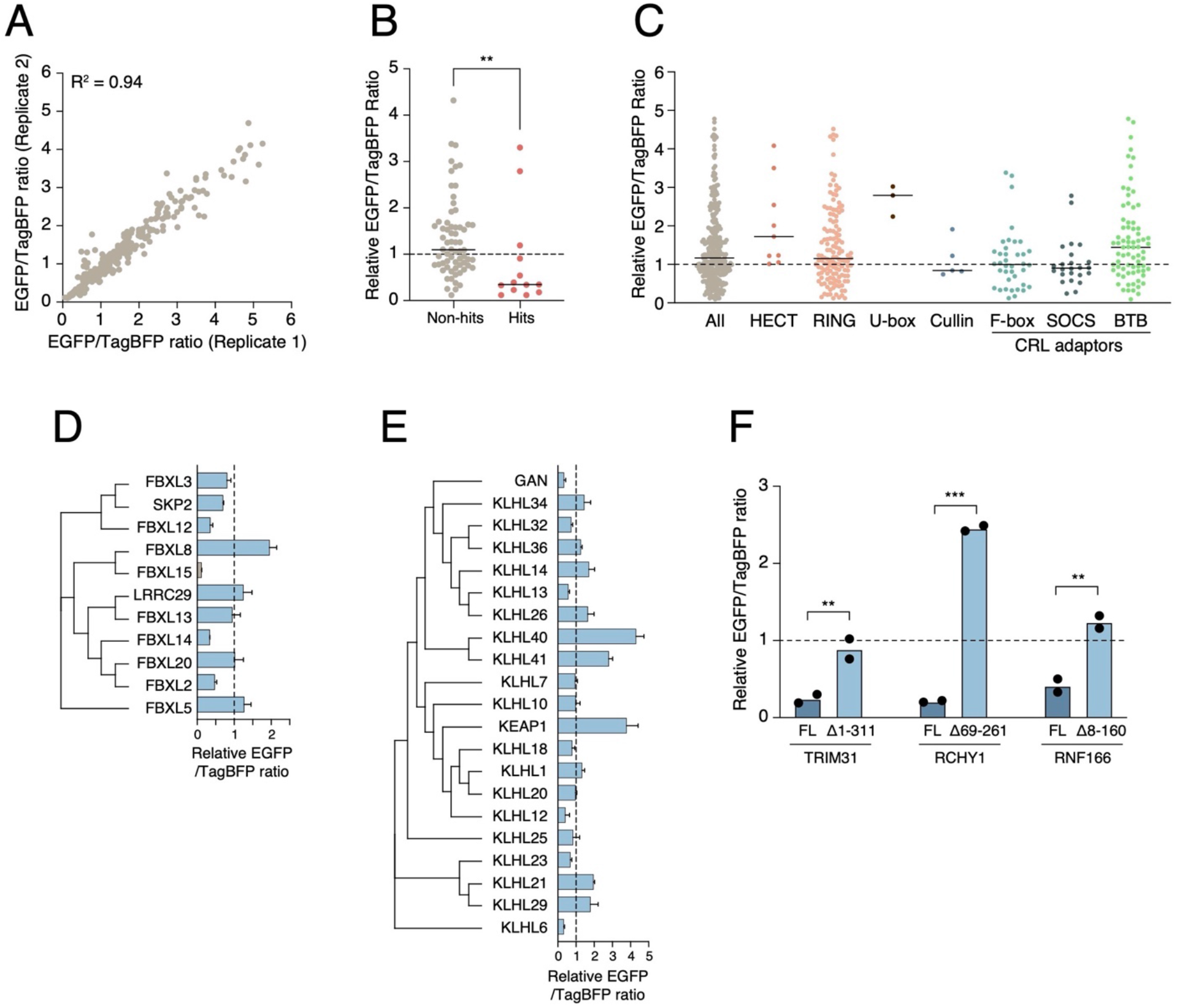
Analysis of E3 ligase activity in proximity-dependent degradation. Related to Figure 2. **(A)** Comparison of two independent replicates for the degradation assay with 290 individual vhhGFP-tagged E3 ligases. (**B**) Comparison of hits recovered from the pooled ORFeome screens and non-hits in arrayed degradation assay. Statistical significance was calculated with unpaired two-tailed Mann Whitney test (**, p < 0.01). (**C**) Effect of different E3 ligase families on EGFP stability. HECT, Homologous to E6AP C terminus; RING, Really Interesting New Gene; BTB (also known as POZ), BR-C, Ttk and Bab; CRL, Cullin E3 RING ubiquitin ligase. (**D**) Phylogenetic clustering of FBXL family E3 ligases and their activity in the degradation assay. (**E**) As in D, but for BTB-BACK-Kelch family E3 ligases. (F) Comparison of the activity of full-length E3 ligase constructs to their splice variants without the RING domain. FL, full length. Statistical significance was calculated with an unpaired two-tailed t test assuming equal variance (**, p < 0.01; ***, p < 0.001).

**Figure S4.**
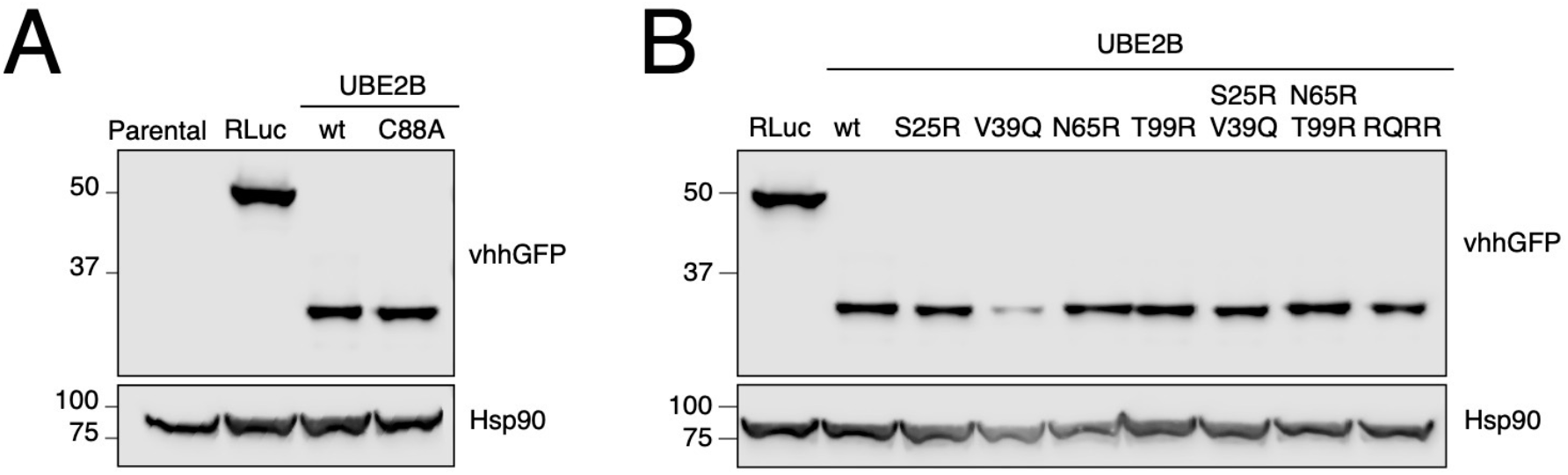
Expression of UBE2B mutant constructs. Related to Figure 3. **(A)** Western blot analysis of wild-type UBE2B and the catalytically inactive mutant C88A. (B) Expression of UBE2B mutants deficient in E3 binding. RQRR, S25R/V39Q/N65R/T99R quadruple mutant.

**Figure S5.**
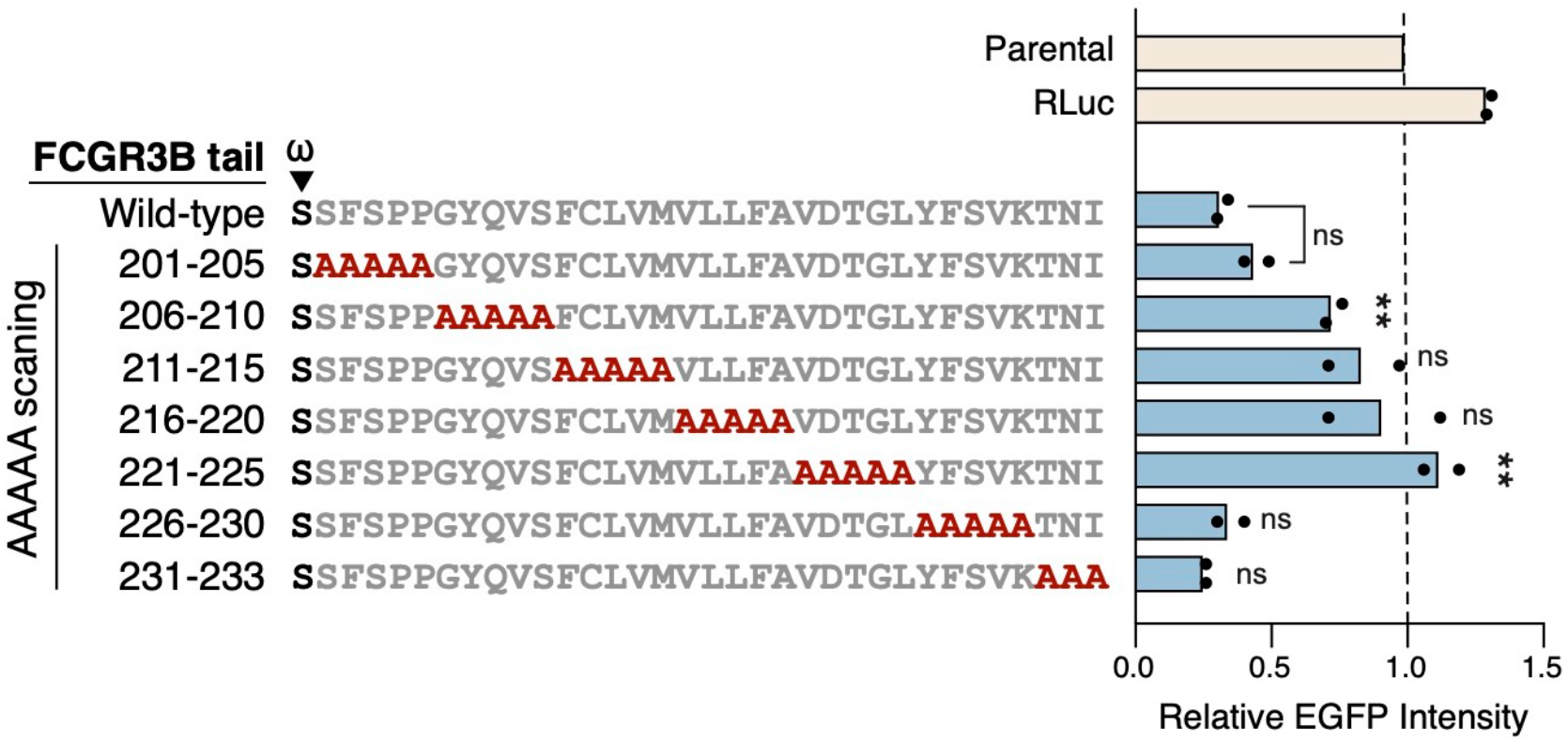
Alanine scanning of the C-terminal tail of FCGR3B. Related to Figure 4. Indicated FCGR3B C-terminal tail mutants fused to vhhGFP were assessed in the EGFP-ABI1 degradation assay. Statistical significance was calculated with one-way ANOVA with Dunnett’s correction for multiple hypothesis.

**Figure S6.**
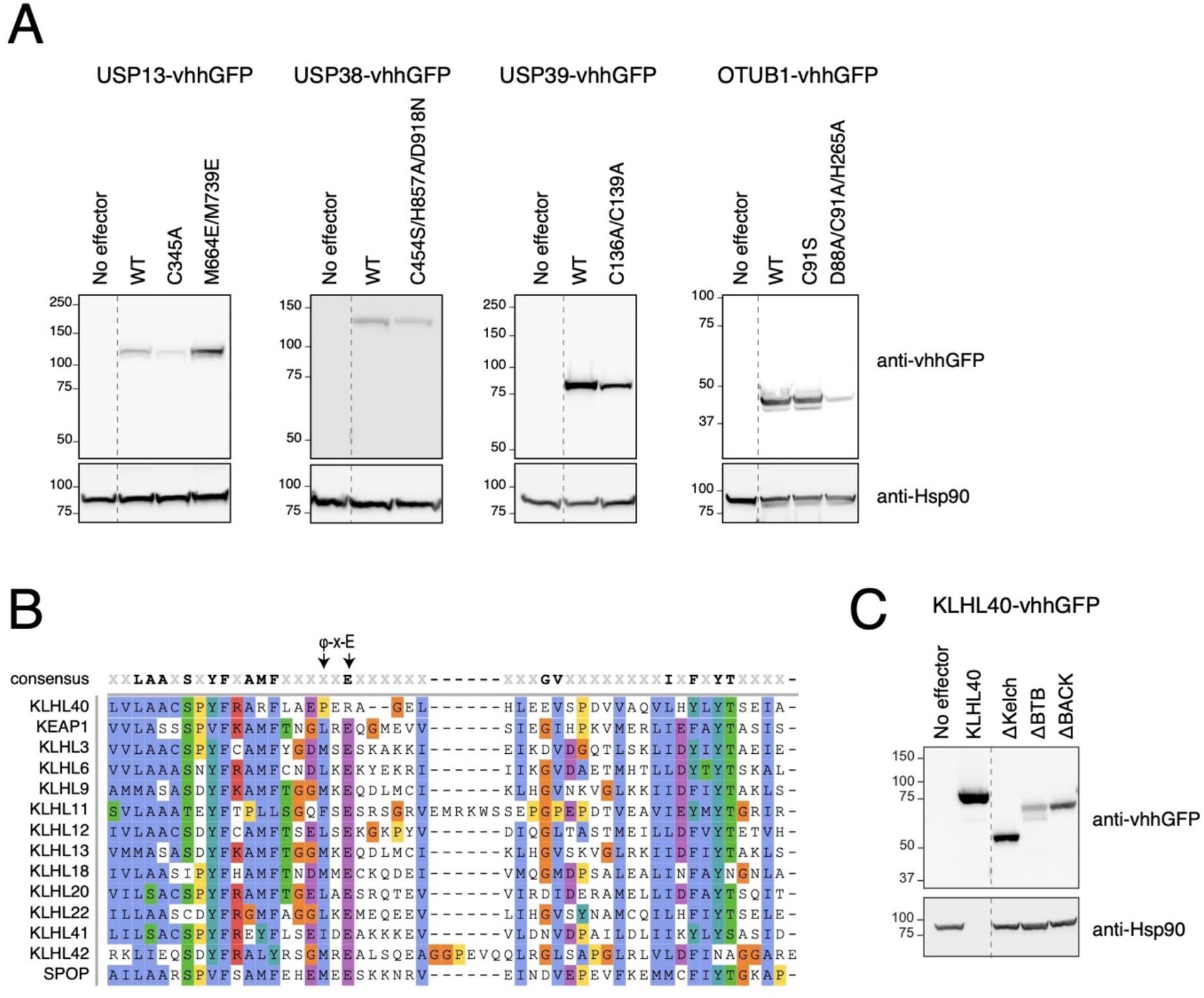
Deubiquitinases and KLHL40 as proximity-dependent stabilizers. Related to Figure 5. (**A**) Western blot analysis of deubiquitinase mutants fused to vhhGFP transfected into 293T cells. (**B**) Multiple sequence alignment of BTB domains from different BTB-BACK-Kelch domains proteins and SPOP (binds CUL3), colored by conservation. The φ-X-E motif residues (where φ represents a hydrophobic amino acid) that binds CUL3 is annotated with arrows. (**C**) Western blot analysis of KLHL40 constructs fused to vhhGFP transfected into 293T cells. Note that Hsp90 band could not be observed on the KLHL40 lane after stripping the blot; however, the same amount of total lysate was loaded.

**Figure S7.**
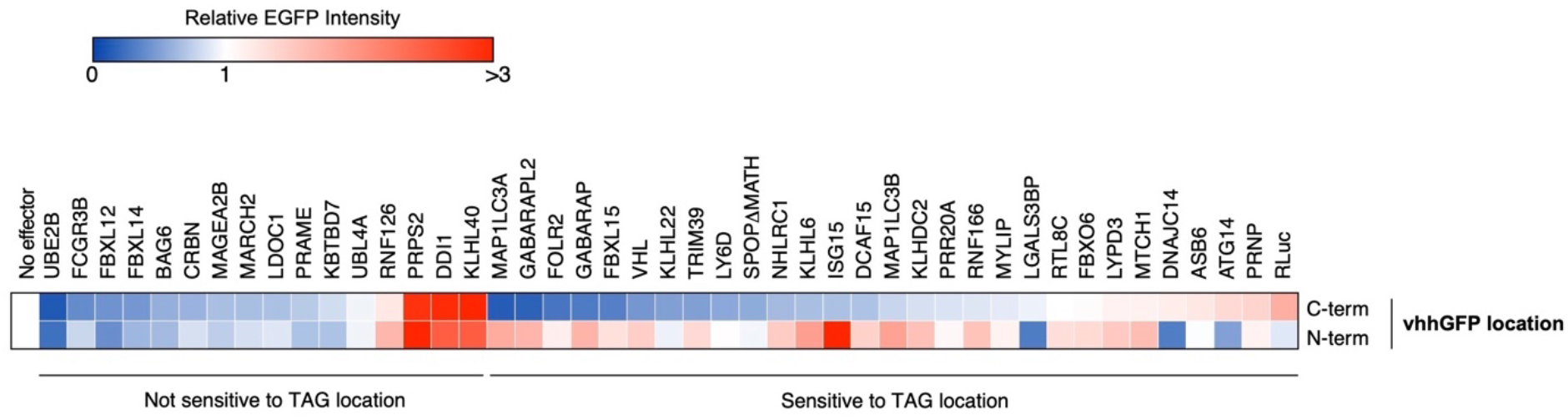
Effect of tag location on effector activity. Related to Figure 6. The EGFP-ABI1 293T reporter cell line was transfected with indicated effectors fused to vhhGFP in their C terminus or N terminus. EGFP fluorescence was measured by flow cytometry and normalized to cells transfected with an unrelated construct.

**Figure S8.**
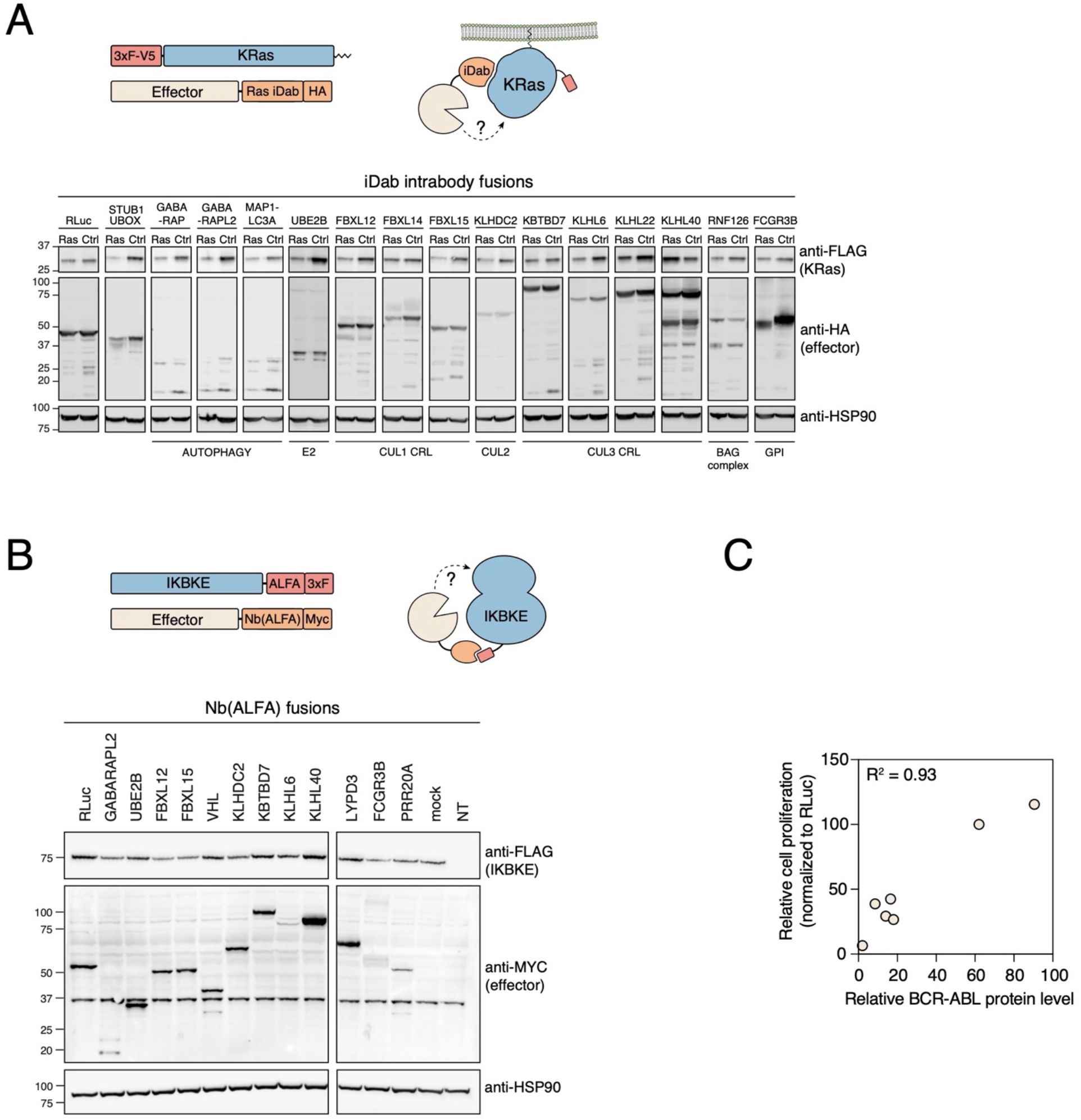
Targeting non-GFP tagged proteins with novel effectors. (**A**) HeLa cells were co-transfected with 3xFLAG-V5-KRAS and indicated effectors fused to an intracellular single domain antibody (iDab) targeting Ras or LMO2 (control). (**B**) 293T cells were co-transfected with IKBKE fused to ALFA-3xFLAG tag and indicated effectors fused to Nb(ALFA)-Myc. (**C**) Correlation between K562 cell proliferation and BCR-ABL levels in cells stably expressing indicated effectors fused to Mb(ABL) treated with doxycycline for 6 days.

## METHODS

### Cell lines

HeLa Kyoto, HCT116, and all 293T cells, including the EGFP-ABI1-IRES-TagBFP reporter cell line used for screens, were maintained in DMEM supplemented with 10% fetal bovine serum (FBS) and 1% penicillin-streptomycin. K562 cells were cultured in RPMI supplemented with 10% fetal bovine serum (FBS) and 1% penicillin-streptomycin. 293T, K562, and HCT116 Tet-inducible cell lines were maintained in their respective regular media supplemented with 10% Tet system approved FBS (Gibco A47363-01) and 1% penicillin-streptomycin. Cells were maintained at 37°C in a humidified incubator at 5% CO_2_ and routinely tested for mycoplasma contamination.

### Plasmids

Unstable mutant targets were cloned into pcDNA3.1-[ORF]-GSlinker-EGFP-P2A-DsRed destination vector, using Gateway cloning technology. For the vhhGFP and PYL1 degradation assays, entry clones were picked from the hORFeome collection and subcloned into pcDNA3.1-[ORF]-GSlinker-vhhGFP-SV40-TagBFP, pcDNA3.1-[ORF]-GSlinker-vhhGFP, pcDNA3.1-vhhGFP-ORF and pcDNA3.1-[ORF]-GSlinker-PYL1 destination vectors. For WDR5 endogenous protein degradation, effector-coding sequences were cloned into pSTV6-TetO-[ORF]-Mb(S4) WDR5-HA lentiviral plasmid, allowing expression of the respective proteins with a C-terminal monobody Mb(S4) recognizing WDR5 with a high affinity (Gupta et al., 2018). For BCR-ABL endogenous protein degradation, effector coding sequences were cloned into pSTV6-TetO-[ORF]-AS25-HA lentiviral vector, allowing expression of the respective proteins with a C-terminal high affinity monobody AS25 directed to the Src homology 2 (SH2)-kinase domain interaction interface (Wojcik et al., 2016). To generate ALFA tagged ARAF or IKBKE, their open reading frames were cloned into the Gateway-compatible pcDNA3.4-ALFA-3xFLAG-ORF and pcDNA3.4-[ORF]-3xFLAG-ALFA vectors, respectively. ALFA nanobody fused effectors were generated by subcloning each effector gene into the Gateway compatible pcDNA3.4-[ORF]-NbALFA-Myc plasmid.

For ectopic K-Ras expression, KRAS cDNA was cloned into pcDNA3.1-3xFLAG-ORF destination vector. The effector coding sequences were cloned into pcDNA3.4-[ORF]-Ras iDab-HA and pcDNA3.1-[ORF]-LMO2 iDab-HA vectors, allowing expression of the respective proteins with a C-terminal monobody recognizing K-Ras or a control protein (LMO2), respectively (Tanaka et al., 2007, 2011).

Point mutants were generated by site-directed mutagenesis and deletion constructs were generated by PCR.

### Lentivirus production

Lentiviral particles containing the pooled ORFeome were produced by transfecting HEK-293T cells with pLX301-[ORF]-PYL1 or pLX301-[ORF]-vhhGFP, psPAX2 (Addgene #12260) and pVSV-G (Addgene #8454) vectors at a ratio of 8:8:1. Transfection was performed using Lipofectamine 2000 (Thermo Fisher Scientific, 11668019) on 15-cm dishes according to the manufacturer’s protocol. The medium was changed 24 hours post-transfection. 72 hours after transfection, supernatant was filtered (0.45 μM), pooled, and collected. A similar protocol was followed for small scale virus production when establishing individual stable cell lines with transfection being performed on 6-well plates using Lipofectamine 2000 reagent.

### Stable cell line generation

A monoclonal 293T cell line expressing EGFP-ABI1-IRES-TagBFP was generated by sorting single cells by FACS after lentiviral infection and blasticidin (6 μg/ml) selection. A clone showing high EGFP and TagBFP expression was selected for subsequent experiments.

To generate the inducible doxycycline-inducible cell lines, effectors were subcloned into Gateway compatible pSTV6-TetO-[ORF]-EGFP, pSTV6-TetO-[ORF]-AS25-HA or pSTV6-TetO-[ORF]-Mb(S4) WDR5-HA lentiviral plasmids. 293T and HCT116 cells were infected in the presence of 8μg/mL polybrene and selected with 3 μg/ml puromycin 24 hours post infection. K562 cells were infected by spin-down at 3000 rpm for 90 minutes in the presence of 8 μg/ml polybrene and selected with 3 μg/ml puromycin 24 hours post infection.

To enrich for high EGFP expression with fusion proteins, cells infected with pSTV6-TetO-[ORF]-EGFP lentivirus were induced with 1 µg/ml doxycycline and sorted for high EGFP population.

### Pooled ORFeome library generation

Entry clones from the human ORFeome collection (v8.1) were collected into 40 standardized subpools each containing ∼384 ORFs and cloned into the lentiviral Gateway-compatible destination vector pLX301-[ORF]-PYL1 or pLX301-[ORF]-vhhGFP. LR reactions were set up in duplicates with 150 ng of each entry ORF subpool, combined with 1 μl of Gateway LR clonase II in a total of 5 μl reaction volume and incubated overnight in TE buffer at room temperature. For the next two days, 1 μl additional LR enzyme in 4 μl TE and 150ng destination vector was added to each reaction. Subpools were transformed into a chemically competent Stbl3 *E. coli* strain and spread on LB agar plates containing ampicillin (100 μg/μl) overnight at 30°C. Colonies were counted to ensure >200-fold coverage, collected in SOC on ice, pelleted and maxiprepped on multiple columns based on weight of the dry pellets.

### Pooled screens

ORFeome libraries tagged at the C-terminus with PYL1 or vhhGFP were packaged into lentiviral particles. The EGFP-ABI1-IRES-TagBFP reporter cell line was transduced at low multiplicity of infection (MOI) with approximately 30% cell survival after puromycin (1 μg/mL) selection. Untransduced cells under the same condition were fully eliminated. Sufficient cells were transduced to maintain >500-fold coverage of the libraries. For the ORFeome-PYL1 library, recruitment was induced by treating cells with 100 μM abscisic acid (ABA, Sigma) for 48 hours. In parallel, a control batch of cells were treated with equal volume of DMSO. Cells were then washed in 1 x PBS, treated with dissociation buffer (1 mM EDTA, 10 mM KCl, 150 mM NaCl, 5 mM sodium bicarbonate, 0.1% glucose) and resuspended in flow buffer (5 mM EDTA, 25 mM HEPES pH 7, 1% BSA, PBS). For each library, high EGFP (top 10%) and low EGFP (bottom 10%) populations were sorted (in duplicate or triplicate) using BD FACS Melody (kindly provided by the Stagljar lab, CCBR) or Sony Biotechnology LE-MA900FP Multi-Application cell Sorter (Flow Cytometry Facilities, TCP, Lunenfeld-Tanenbaum Research Institute). Genomic DNA was directly extracted using QIAmp DNA Blood Mini Kit (QIAGEN).

### ORFeome sequencing

Nested PCR was performed using all the purified genomic DNA from sorted populations or at least 5 μg of genomic DNA from unsorted populations. ORFs were amplified from genomic DNA using primers targeting the T7 promoter (CGACTCACTATAGGGAGACCCAAG) and PYL1 (ATTCATCTTGCGTTGGTGCTCC) or vhhGFP (GCCACCAGACTCCACCAGTTGGAC). The product of this reaction was pooled for each sample and further amplified by primers targeting sequences just outside the Gateway attB sites (CAGTGTGGTGGAATTCTGCAG and CCGCCACTGTGCTGGATATC) for an additional 10 cycles. Amplicons were subsequently separated on 1% agarose gel and any visible PCR product excluding primer dimers were gel purified. After quantifying DNA using the Quant-iT 1X dsDNA HS kit (Thermo Fisher Scientific, Q33232), 50 ng per sample was processed using the Illumina DNA Prep, (M) Tagmentation kit (Illumina, 20018705), with 6 cycles of amplification. 2 μl of each purified final library was run on an Agilent TapeStation HS D1000 ScreenTape (Agilent Technologies, 5067-5584). The libraries were quantified using the Quant-iT 1X dsDNA HS kit (Thermo Fisher Scientific, Q33232) and pooled at equimolar ratios after size-adjustment. The final pool was quantified using NEBNext Library Quant Kit for Illumina (New England Biolabs, E7630L) and paired-end sequenced on an Illumina MiSeq.

### Analysis of sequencing data from pooled activation screens

An index of the ORFeome reference sequences was created using the STAR aligner v2.7.8a. Reads from the ORFeome libraries were aligned with the STAR aligner allowing a maximum of 3 mismatches. To identify degradation and stabilization effectors, the edgeR package (Robinson et al., 2010) was used to calculate log2 fold change, p-value, and false discovery rate (FDR) for each ORF by comparing changes in counts from sorted samples to unsorted cells.

### vhhGFP degradation assay for individual effectors

Degradation assays with individual effectors were performed in a 48-well cell culture format EGFP-ABI1-IRES-TagBFP reporter cell line was transfected using Lipofectamine 2000 (Life Technologies) with 200 ng of the effector fused to vhhGFP or PYL1 and 15 ng of transfection control plasmid expressing 3xFLAG-tagged DsRed. For the unstable mutant stabilization experiments, 293T cells were co-transfected with 100 ng of the effector fused to vhhGFP and 100 ng of the target fused to EGFP. For the effector-PYL1 constructs, recruitment was induced by treating cells with 100 μM abscisic acid (ABA, Sigma) for 48 hours. In parallel, a control batch of cells were treated with equal total volume of DMSO. 48 hours post-transfection, cells were washed in 1 x PBS, treated with dissociation buffer, and resuspended in flow buffer. Cells were spun down in a microcentrifuge at 1000 rpm for 5 minutes. Cell pellets were resuspended in flow buffer and analyzed using BD LSR Fortessa or BD LSR Fortessa X20 (BD Biosciences; University of Toronto Faculty of Medicine Flow Cytometry Facility).

### K-Ras degradation assay

HeLa cells were seeded in 12-well plates (100,000 cells/well) and transfected 24 hours later with 200 ng of 3xFLAG-KRAS and 400 ng of effector fused to Ras iDab or LMO2 iDab 48h hours after transfection, cells were harvested and subjected to western blot analysis.

### Endogenous protein degradation assay

K562 and HCT116 stable cell lines were seeded in 12-well plates (1 x 10^6^ cells/well). K562 cells were incubated with 1 µg/ml of doxycycline or 1% DMSO for 48 hours. HCT116 cells were incubated for 24 hours to let the cells acclimatize before being treated with 1 µg/ml of doxycycline or 1% DMSO for another 24 hours. Cells were then harvested and subjected to western blot analysis.

### ALFA-tagged kinase degradation assay

ALFA-tagged kinase degradation assays were performed in a 12-well cell culture format (100,000 cells/well). After 24 hours, 293T cells were transiently transfected with 1 µg of either pcDNA3.4-ALFA-3xFLAG-TEV-ARAF or pcDNA3.4-IKBKE-3xFLAG-ALFA, using Lipofectamine 2000 (Life Technologies) following the manufacturer’s instructions. Cells were harvested and subjected to western blot analysis 16 hours post-transfection.

### Western blot

Cells were lysed in CSK lysis buffer (20 mM Hepes-KOH pH 7.9, 100 mM NaCl, 1 mM MgCl_2_, 1 mM EDTA, 300 mM sucrose, 1 mM DTT, 0.1% Triton X-100, benzonase, and protease inhibitor cocktail). For BCR-ABL, WDR5, ARAF and IKBKE degradation assay, cells were lysed in NP40 lysis buffer (50 mM Tris-HCl pH 7.6, 150 mM NaCl, 1% NP40 and protease inhibitor cocktail). After centrifugation at 16,000 g for 5 min at 4°C, the same amount of each cellular lysate was analyzed by gel electrophoresis and western blot using anti-WDR5 antibody (D9E1I; Cell Signaling #13105), anti-HSP90 antibody (F-8; Santa Cruz Biotechnology), anti-HA (Sigma #H6908), anti-c-ABL (Cell Signaling #2862T), anti-MYC (BioLegend #626802), and anti-FLAG (DSHB #12C6c) as the primary antibody. Goat HRP-conjugated anti-rabbit IgG (Cell Signaling #7074S) or anti-mouse IgG (Cell Signaling #7076S) were used as secondary antibodies. MonoRabTM HRP Rabbit anti-Camelid VHH antibody (GenScript #A01861) was used to detect vhhGFP fusion proteins. Chemiluminescence signal was generated with Immobilon Western Chemiluminescent HRP Substrate (Millipore) and detected with MicroChemi4.2 (FroggaBio).

### Inhibitor treatments

Cells were treated with 1 µM MLN4924 (Chemietek) for 24 hours, 100 nM Bortezomib (Calbiochem) for 6 hours, 20 µM cycloheximide (Sigma) for 6 hours, 2.5 µM CB-5083 (Selleck) for 6 hours, or 0.01% DMSO (Fisher bioreagents) for 24 hours.

### Proliferation assay of stable K562 cell lines

Proliferation assay was performed in a 96-well culture format (1,000 cells/well). Cells were grown in the presence of 1 µg/ml doxycycline or 1% DMSO for the indicated times. Doxycycline or DMSO were newly added after 72 hours. CellTiter Glo (Promega) was used to measure cell viability, following the manufacturer’s instructions. Luminescence intensities were measured using a multimode microplate reader (Biotek).

### Immunofluorescence

HeLa Kyoto cell were seeded into opaque black, clear bottom 96-well plates at 4,500 – 5,000 cells per well. The next day, cells were transfected using XtremeGENE 9 (Roche), as per the manufacturer’s instructions. 48 hours after transfection, cells were washed with 1 x PBS and then fixed for 15 minutes at room temperature with 4% paraformaldehyde in DMEM containing 10% FBS. Following fixation, cells were washed three times in 1 x PBS, permeabilized with 0.1% Triton X-100/1 x PBS, and then blocked with blocking buffer (0.1% Triton X-100/1 x PBS/1% BSA) for 30 minutes at room temperature. After blocking, fixed cells were incubated with MonoRab^TM^ iFluor 647 Rabbit Anti-Camelid VHH antibody (GenScript A01994) and Hoechst 33342 diluted in blocking buffer for 1 hour at room temperature. Finally, cells were washed three times in 1 x PBS and imaged using the Opera Phenix high-content microscope (Perkin Elmer) at 63x magnification.

